# Factorization and spatial encodings: a hypothesis about the foundations of the genomic code

**DOI:** 10.64898/2026.07.14.738413

**Authors:** Karim G. Habashy, Benjamin D. Evans, Dan F. M. Goodman, Jeffrey S. Bowers

## Abstract

The genomic mechanisms that efficiently encode the initial architecture and synaptic connectivity of neural circuits remain poorly understood. We hypothesise that two primary mechanisms — spatial encoding and factorisation — enable a limited genome to initialise networks of billions of neurons. Spatial encoding, a form of indirect representation, compresses neural network parameters while enforcing structural continuity. Complementarily, we introduce a factorisation mechanism inspired by reaction-diffusion models, comprising a spatially invariant reaction rule and spatially variant diffusion dynamics. Based on this insight, we can efficiently abstract a neural network layer as a spatially invariant weight kernel (or filter) and its spatially variant transformations along the spatial dimensions. Thus, coupling this variant-invariant decomposition with spatial encodings can substantially reduce the size of the solution space explored by the genome. In addition, we show that this coupling leads to efficient itialisation of cortical maps, such as V1 orientation maps, and neural networks with the ability to generalise. In summary, coupling factorisation and spatial encodings can offer functional advantages to the evolving genome.

## 1 Introduction

Unlike the uniform architectures of many artificial neural networks, the genomic initialization of the brain is intrinsically heterogeneous. The brain is anatomically structured across multiple spatial scales, manifested as distinct neuronal types at the microscale and specialized topological architectures at the macroscopic network level. Topographic feature maps (Blasdel & Salama, 1986) are a prominent example of the latter, in which the neuronal receptive fields vary smoothly across space. Representative examples include tonotopic and periodotopic maps in the auditory cortex, alongside orientation tuning and ocular dominance maps in the visual cortex. Regarding visual maps, There is strong consensus that such feature maps can arise innately, independent of sensory experience (Crair et al., 1998; Crowley & Katz, 1999; White et al., 2001), although postnatal experience is required to maintain their long-term responsiveness and selectivity (Crair et al., 1998; Cloherty et al., 2016).

Beyond the structural scaffolding of feature maps, the genome incorporates diverse evolutionary biases during brain initialization. For instance, neonates — both human and avian — exhibit an innate preference for face-like stimuli immediately following birth or hatching (Salva et al., 2011; Kobylkov et al., 2024). This predisposition represents a primitive form of inductive bias as it can be defined by a consistent response to novel, *out-of-distribution*, stimuli after birth without postnatal learning. In general, it has been argued that such aspects of generalization are inherently innate (Ghirlanda & Enquist, 2003; Wang et al., 2024).

Despite the ubiquity of these innate biases, only feature maps have been extensively studied from a mechanistic perspective in an in-life learning framework. Traditional models often rely on activity-dependent self-organizing principles (Durbin & Mitchison, 1990; Kaschube et al., 2010) emphasizing input selectivity coupled with center-surround (local excitation/global inhibition) dynamics (Bednar, 2012; Stevens et al., 2013). However, there is currently no account of how these maps emerge via indirect encoding, where a genotype generates the structure (like weights) of a phenotype network. Similarly, the role of genomic generalizations (or inductive biases) has not been systematically investigated through the lens of indirect representations. Here, we demonstrate that these innate biases can emerge efficiently from a minimal set of tunable parameters rooted in the same mechanistic principles: weight factorization and spatial encoding, which is a form of indirect encoding.

The field of indirect encoding was motivated by the genomic developmental program (Stanley, 2007). Thus, work on indirect encoding deals with encoding a function (the phenotype) by another function (the genotype). Both the genotype and the phenotype can be modelled with neural networks, where the former generates/encodes the weights of the latter. When the genotype encodes the phenotypic weights as a function of the neuronal coordinates (Fig. 1), we call it spatial encodings. This formulation enforces structural continuity, ensuring that proximal neurons possess similar synaptic weights — a framework pioneered by Compositional Pattern Producing Networks (CPPNs; Stanley, 2007; Stanley et al., 2009).

**Fig. 1:**
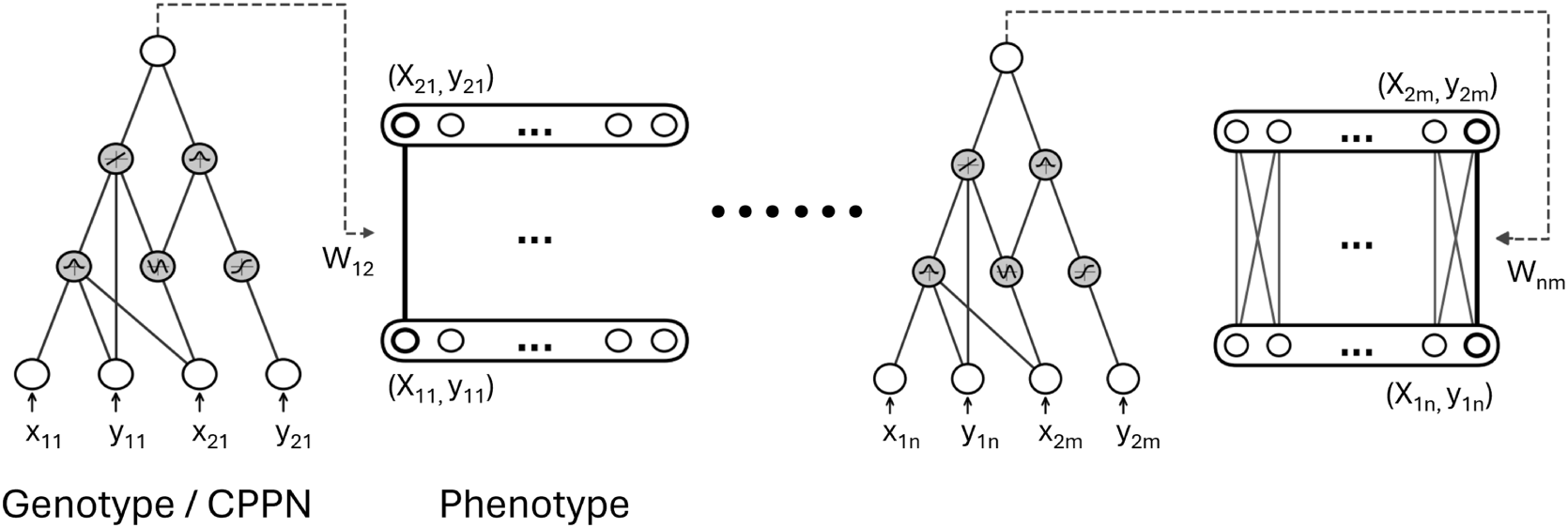
Spatial encodings as a model of the genomic developmental program. A compositional pattern producing network (CPPN) takes in as an input the coordinate pair of pre- and postsynaptic neurons and outputs their associated weights. A CPPN utilizes, in its hidden layer, various activation functions that assist it in mimicking some aspects of the genomic encoding, like producing weight structures with bilateral symmetries (Gaussian functions) and repetitions (Sinusoidal functions). This figure is inspired by Sosa & Stanley, 2018.

Here we show that spatial encodings are sufficient to produce various types of innate structures, like V1 orientation maps. This provides a novel account of V1 organization compared to previous work that relied on in-life learning algorithms (Bednar, 2012; Philips & Chakravarthy, 2017). However, these V1 maps can emerge more efficiently, and at a better performance level, when combing spatial encodings with factorization during in-life learning. The performance level is measured by the average final training loss after 2000 epochs. In machine learning, weight factorization typically refers to low-rank approximations or Singular Value Decomposition (SVD) used for post-training compression (Novikov et al., 2015; Swaminathan et al., 2020), regularization (Gunasekar et al., 2017), or parameter-efficient fine-tuning (Hu et al., 2021). We propose a novel reinterpretation of factorization inspired by reaction-diffusion dynamics: neural weights are decomposed into a spatially invariant kernel and its transformations across spatial dimensions. While this formalism resembles the generation of filter banks or dictionaries in sparse coding literature (Olshausen & Field, 1997; Kavukcuoglu et al., 2010), it differs fundamentally; whereas standard dictionaries assume linearly independent, uncorrelated kernels, our framework produces kernels that are intrinsically correlated through spatial transformations. This kind of factorization can prove useful for an evolving genome as it reduces the size of the parameter search space.

In summary, we provide a theory of the genomic code (the code defined by a set of genes) by integrating the functional properties of weight factorization with spatial encodings. Using this integrated framework, we demonstrate i) efficient encoding of orientation maps, ii) efficient initialization of generalizable features and iii) efficient encoding of arbitrary innate functions as exemplified by the MNIST dataset (LeCun et al., 2002).

## 2 Methods

This section begins with a presentation of the phenotypic hidden layer structure used in this work, which differs from conventional convolutional neural networks. We then introduce *differentiable* CPPNs where the term differentiable is used to disambiguate them from conventional CPPNs which use evolution for training. Next, we demonstrate our factorization approach, inspired by Jaderberg et al. (2015). Finally, we present the various architectures combined with the kernels used in this work.

### 2.1 Phenotypic hidden layer architecture

The layout used in this work is influenced by the 2D horizontal arrangement of feature selectivity in V1 (Fig. 2a,i). This kind of arrangement can be contrasted with the vertical (stacked) arrangement of feature maps in depth-wise CNNs (Fig. 2a,ii). In terms of feature extraction in the first hidden layer, both layouts can be argued to work in the same way, i.e, they can extract similar features. However horizontal arrangement can have two advantages: i) a biological advantage: CNNs separate features into different maps, which does not appear to be the case in the brain; ii) a structural advantage: CNNs treat feature maps as independent and uncorrelated within the same stack/layer. For example, a 120*^o^* line orientation feature map can be positioned between distant orientation maps like 0*^o^* and 40*^o^*. CNN kernels do not enforce ordering or continuity. Conversely with horizontal arrangement, and under the constraint of continuity, nearby orientations are spatially proximal on the same map.

**Fig. 2:**
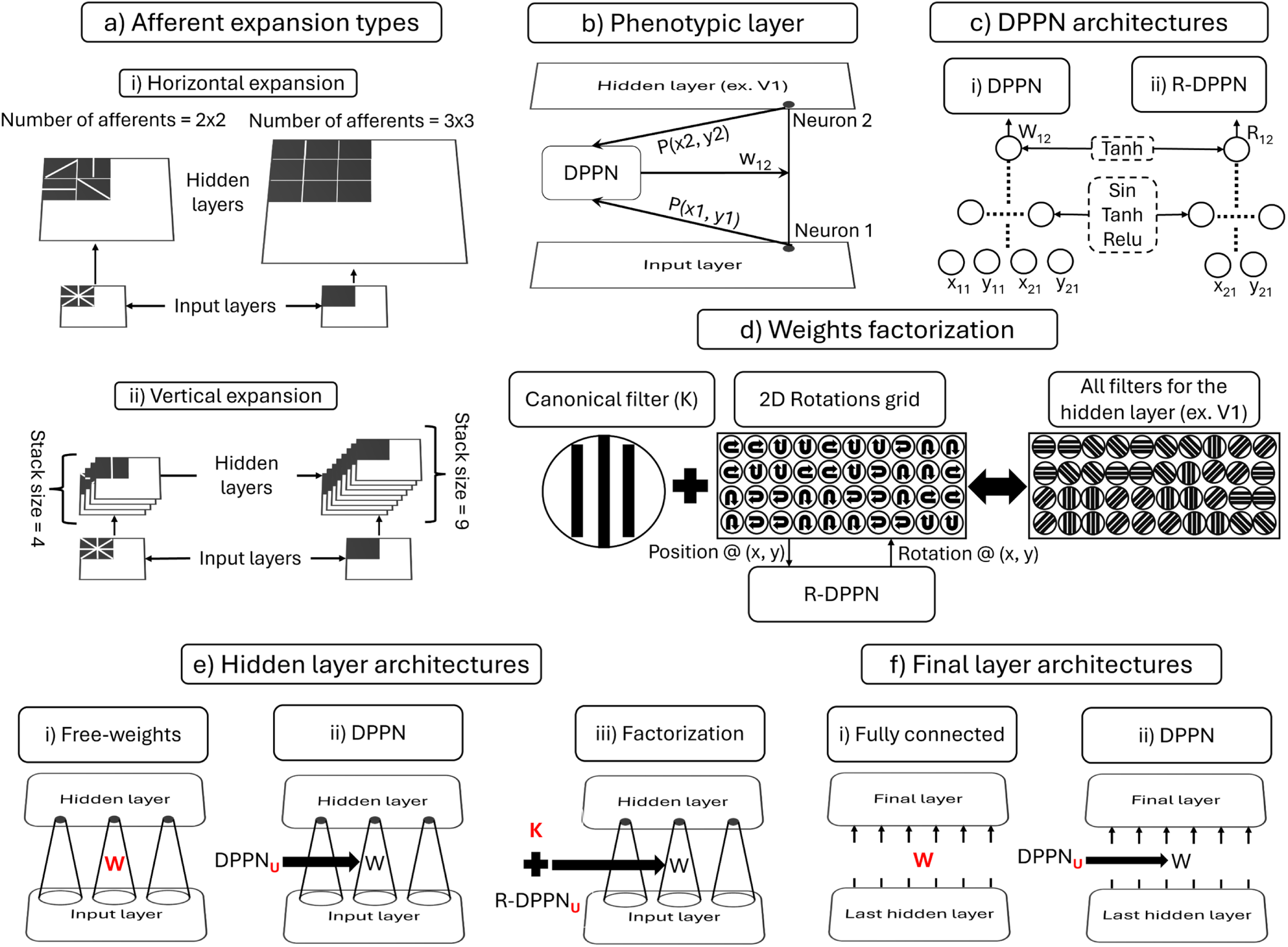
Employed models and network architectures. **a)** afferent expansion: how many times a patch in an input layer is copied and replicated into the next layer to be convolved with. There are two modes of expansion, horizontal, which is used in this work, and vertical, which is the traditional approach in depth-wise convolutional neural networks. **b)** genomic encoding of phenotypic weights: a differentiable pattern producing network (DPPN) takes in the spatial locations of pre- and postsynaptic neurons respectively, and outputs the associated weight. **c)** types of DPPNs used in this work: a DPPN is used to encode weights, while a rotations-DPPN (R-DPPN) is used to encode the rotations of the canonical weight kernel. Also shown are the activation functions used in the final and hidden layers. **d)** weight factorization: a hidden layer can be composed of a kernel (canonical filter) and its rotations along the layer’s spatial dimensions, effectively giving us the complete kernels for a hidden layer. The 2D map of rotation angles can be parametrized by a 2D array, or encoded by an R-DPPN. **e)** hidden layer architectures: i) free-weights, where the weights are directly learnable. ii) DPPN, where the hidden weights are encoded by a DPPN. iii) factorization, where the hidden weights are learned by a canonical filter and its rotations along the spatial dimensions of the layer. **f)** final layer architectures: i) fully connected, where every output neuron is connected to every neuron in the previous layer. ii) DPPN, where the two layers are fully connected but their weights encoded with a DPPN. Red symbols are the learnable parameters with **W** symbolizing the parameters of the phenotype and **U** the parameters of the genotype.

### 2.2 Differentiable CPPNs

In machine learning, the conventional approach to learning functional mappings is to use a neural network that is mainly parametrized by neuronal weights. In this approach, the weights are adjusted (or fine tuned) to achieve a desired input-output mapping. We call this approach *direct encoding* as the weights are directly adjusted to learn the function or mapping. Conversely, another approach to specifying the weights of the main neural network, which we call the *phenotype*, is to redefine them as the output of another neural network, which we call the *genotype*. In this formulation, the phenotypic-weights are not learned, but are parametrized by the genotype network, which are specified by its learned genotypic-weights - an approach we call *indirect encoding*.

Compositional pattern producing networks (CPPNs) are a subtype of indirect encoding networks which encode the parameters of another neural network (Stanley, 2007) and are of particular interest to this work. We treat the CPPN as the genotype and the encoded network as the phenotype (Fig. 2b). Unlike other indirect encoding techniques, CPPNs uniquely account for the spatial arrangement of neurons in the phenotypic layers by outputting weights as a function of the relative positions of pre- and postsynaptic neurons (Fig. 2ci). Generally, CPPNs need not only encode the weights of a neural network, they can also be used to generate the values of any scalar function in R as a function of spatial coordinates (*x, y, z, …*), for a 2D example, see (Fig. 2c,ii).

The conventional approach to adapt (or train) CPPNs is by evolving its structure and parameters, like weights and biases (Stanley et al., 2009). However, in this work, we utilize a modified version of CPPNs, with static structure, but differentiable weights and biases, termed *differentiable* pattern producing networks (Fernando et al., 2016) or DPPNs in short. When DPPNs encode weights we use the notation DPPN, when encoding a 2D function, such as a rotation (see weight factorization), we use the notation R-DPPN. The above description can be summarized in (1) and (2), where we contrast the conventional approach of learning neural networks, with our approach of using DPPNs for a single layer.

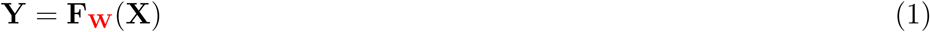

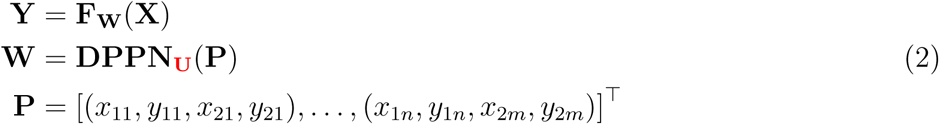

In the above equations, the learnable parameters are highlighted in red. In (1), we show the conventional approach, where a phenotypic function **F**(.), parameterized by learnable phenotypic-weights **W**, maps inputs **X** to outputs **Y**. Conversely, in (2), the weights **W** are not learnable, but are the output of a genotypic function **DPPN**(.) which is parametrized by the learnable genotypic-weights **U**. In machine learning, networks that generates weights for another network are sometimes referred to as hypernetworks (Ha et al., 2016). The function **DPPN**(.) takes as input the positions matrix **P**. The positions matrix is of shape (*mn,* 4), where *m* and *n* are the number of neurons in any two successive layers. Regarding the DPPN networks, to motivate the emergence of periodic structures in the phenotype, we use hidden layer activation functions that are combinations of Sinusoidal, Tanh and Relu functions (Fig. 2c).

### 2.3 Weight factorization

As stated before, the weight factorization employed in this work is an abstraction of the reaction diffusion dynamics. Specifically, as the reaction diffusion equations are built from a spatially invariant reaction rule and a spatially dependent diffusion term, a neural network layer can be built from a spatially invariant weight kernel (canonical kernel) and its spatially dependent transformations (in our case rotations) along the dimensions of the layer (Fig. 2d). This approach is implemented by treating the weight kernel as an image that is differentiably transformed along a neural network layer. This formulation can be compactly summarized in (3) for a single layer.

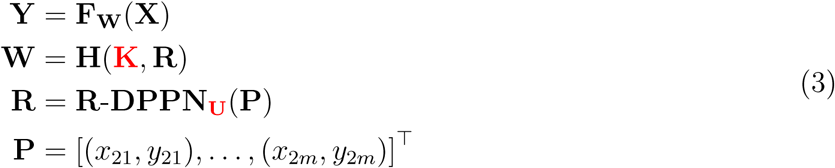

In the above equations, the learnable parameters are highlighted in red. The first line of this equation is the same as (1), except that **W** is not learnable. We acquire **W**, the set of kernels of a layer, by rotating a single learnable kernel **K** along the spatial dimensions of the layer. The rotations at each spatial position are given by the 2D array **R**. It is 2D because we are dealing with only two cartesian coordinates *x* and *y*. The function **H**(.) is a static function that handles the application of the rotations to the canonical filter and the subsequent interpolation. The details of the implementation of **H**(.) can be found in Jaderberg et al. (2015), and it should also be noted that we use bilinear interpolation. Regarding **R**, it can be a freely learnable parameter, or it can be parametrized by a genotypic network **R**-**DPPN**(.), which is parametrized by the learnable genotypic-weights **U**. The function **R**-**DPPN**(.) takes as input the position matrix **P** of shape (*m,* 2), where *m* is the number of neurons in a postsynaptic layer.

### 2.4 Final and hidden layer architectures

One aim of this work is to study the influence of the various layer architectures on the emergent network structure and properties. Thus, we employ a diverse set of hidden and final layer architectures (Fig. 2e and Fig. 2f). For the former, we mainly employ three architectures: i) free-weights, where the weight kernels are free parameters that are directly learned without any restrictions; ii) DPPN, where the weight kernels are indirectly encoded by a DPPN, which enforces continuity; iii) factorization, where the weight kernels are acquired by rotating a canonical filter along spatial dimensions of the layer. In this case, the rotations array is usually indirectly encoded with an R-DPPN. However, in very few cases, the rotations array is directly learned. For the final layer case (Fig. 2f), we show that this layer can be comprised of two architectures: i) fully connected, where every neuron in the final hidden layer is connected to every other neuron in the output layer; ii) DPPN, where the final weights have the same shape (or structure) as the fully connected case, but are acquired through indirect encoding with a DPPN.

## 3 Results

The genome, with its limited information-carrying capacity, can initialize a brain with billions of neurons that comprise a diverse set of functions. For this to be possible, we hypothesize that the genomic code makes use of a constrained set of developmental mechanisms to support this feat. In our work, these mechanisms are limited to factorization and spatial encodings. To demonstrate their utility in initializing diverse brain functions, we begin by showing how V1 orientation maps can efficiently arise from this combination. Subsequently, we demonstrate how this framework explains the efficient emergence of structured innate generalizations. Finally, we illustrate how our model can efficiently encode arbitrary innate functions, exemplified by the widely adopted MNIST benchmark.

### 3.1 V1 orientation maps

Feature maps are regions in the brain where neuronal selectivity (or tuning curves) vary smoothly along the spatial dimensions. These maps are ubiquitous and can be found in sensory areas as well as higher order areas such as the hippocampus where navigational maps (O’Keefe & Dostrovsky, 1971), mediated by place cells, were discovered. In the sensory areas, V1 orientation maps might be argued to be the most exemplary case of such maps. V1 orientation maps are characterized by a slow variation in the orientation tuning along the spatial dimensions. These feature maps are thought to be largely innate, yet there is no account of how such maps can arise from indirect encoding techniques. We demonstrate how the genome can indirectly encode such maps. In addition, we also show how such maps can arise efficiently by combining spatial encodings with factorization.

To support the claims made above, we use a collection of different model network configurations (Fig. 3a), solving a task where the inputs are straight lines with varying orientation, position, thickness and length, while the output is the orientation angle (invariant to everything else). With this setup, we find the emergent V1 orientation map to be the optimal orientation activations of the hidden layer after training.

**Fig. 3:**
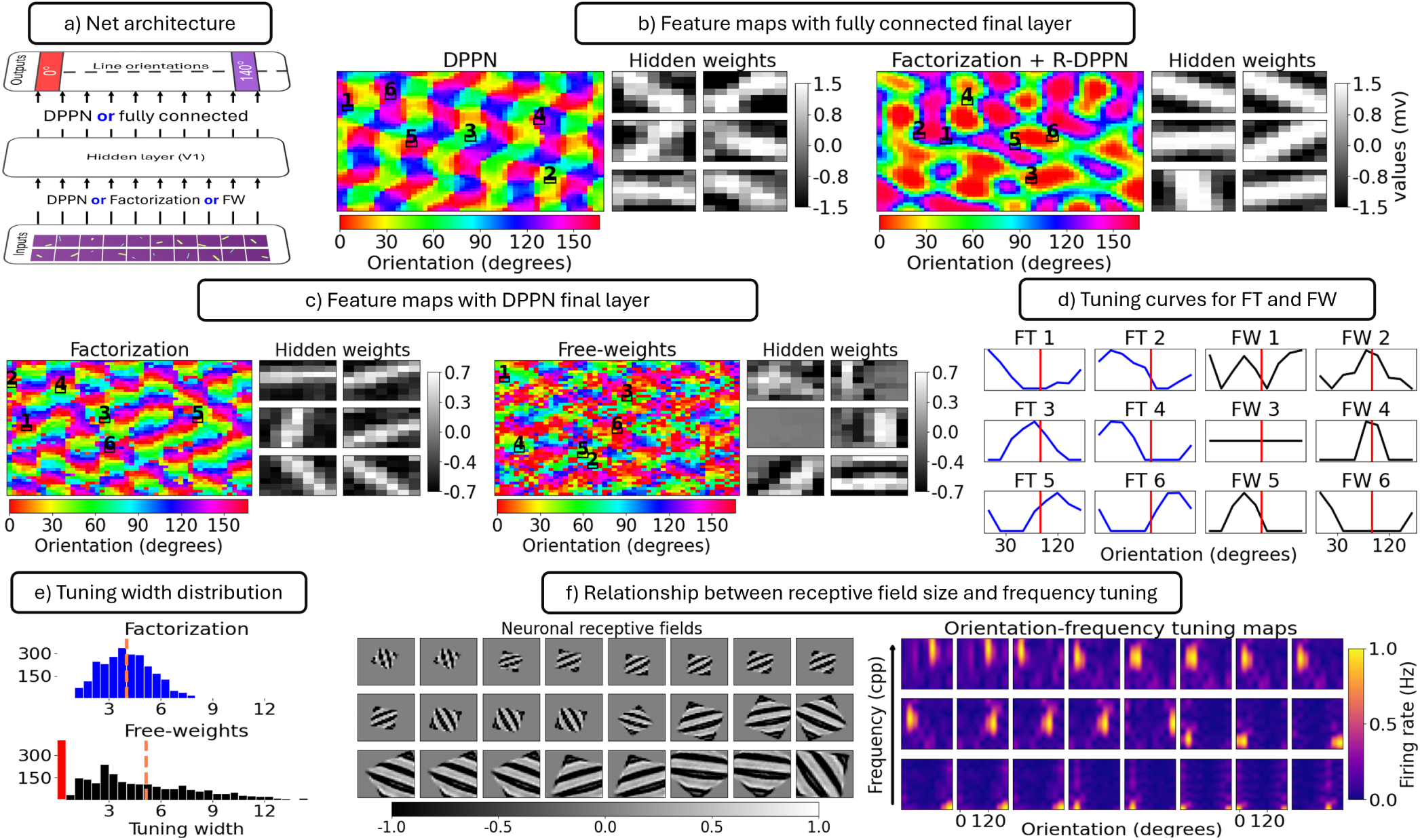
V1 orientation maps. **a)** Network architecture: the final layer can be either a differential pattern producing network (DPPN) or fully connected, while the hidden layer can be a DPPN, free-weights (FW) or factorization. For factorization, continuity can be enforced by a rotations-DPPNS (R-DPPN). **b)** Feature maps with fully connected final layer, while the hidden layer can be a DPPN or factorization + R-DPPN. In this case, continuity is enforced on the hidden layer by the DPPN or the R-DPPN. **c)** feature maps with DPPN final layer, while the hidden layer can be factorization only or free-weights. In these cases, continuity is enforced by the final layer DPPN. **d)** Tuning curves for the selected neurons/kernels in c). The red vertical line marks the 90*^o^* orientation. **e)** Tuning curves width distribution for the left and right maps in c). The orange dashed lines are the averages, while the red bar represents the count of inactive (dead) neurons. **f)** Relationship between receptive field size and frequency tuning: showing on the left example receptive fields with varying sizes, while on the right we show the rate maps of the associated conjunctive tuning of orientation and frequency. For the receptive fields, the gray surround of the resultant kernels arises from the zero padding we applied during the affine transformation to highlight the contribution of the scaling operation.

For the different network configurations, we use different weight architectures for the final and hidden layer to explore their influence on the emergent map structures. Thus, we parameterize the final layer either by a differentiable pattern producing network (DPPN, a network that converts spatial positions into weights), or a fully connected layer. The hidden layer is parameterized by a DPPN, free-weights (i.e. directly learning the weights) or factorization. In the case of factorization, we consider weight kernels that are rotations of a canonical kernel, for example an orientation filter. To rotate this canonical kernel we use a 2 × 3 affine matrix that is parameterized by only one value, the angle of rotation. The 2D scalar map of all rotations of a layer is the rotations map, and it can be trained as a free variable (a 2D array) or encoded by a rotations-DPPN (R-DPPN) to enforce continuity.

In short, we have two variations of the final layer: i) a DPPN, ii) fully-connected, while the hidden layer can be one of four variations: i) a DPPN, ii) free-weights, iii) factorization only, iv) factorization + R-DPPN. Although, many more combinations can be made, these combinations of architectures will be sufficient to highlight the points to be demonstrated in the reminder of this section (see section 2 and section E1 for more details).

#### 3.1.1 Hidden-weight continuity gives rise to orientation maps

Enforcing continuity on the weight structure while training on a line-orientation classification task leads to the emergence of V1-like orientation maps (Fig. 3b left image; see Fig.S1 for more seeds). Continuity was achieved by using a DPPN in the hidden layer, while the output layer was fully-connected. The learned hidden weights in this configuration resemble line filters.

#### 3.1.2 **Efficient map initialization can be achieved by combing factorization and continuity**

Combing factorization with a continuity enforcing algorithm can lead to the emergence of orientation maps (Fig. 3b right image; see Fig. S2 for more seeds). In this case, continuity is applied by indirectly encoding the 2D scalar rotations map by a R-DPPN, which by design, enforces continuity on the rotations values.

Regarding the parameter efficiency of such algorithms, the factorization case (Fig. 3b right image) uses one order of magnitude fewer parameters than the DPPN case (Fig. 3b left image). The DPPN case uses 1020 parameters in the hidden layer, while factorization + R-DPPN only uses 109. In terms of performance, factorization outperforms DPPN by a small margin measured by the mean training loss after 2, 000 epochs: 0.08 ± 0.01 for factorization versus 0.09 ± 0.02 for DPPN. In addition, we performed a comprehensive comparison between factorization and DPPN in terms of parameter count (efficiency), performance (training loss), loss convergence, directionality (sharpness of the oriented kernel structure) and weight structure (Fig.S3, Fig.S4, Fig.S5, Table S1). Finally, we provided a different case of how maps can emerge by combining factorization and a lateral Gaussian kernel in a direct encoding (in-life learning) framework (Fig.S6 and Fig.S7).

#### 3.1.3 Final-weight continuity also gives rise to orientation maps

Swapping the fully connected layer with a DPPN in the final layer leads to the emergence of orientation maps (Fig. 3c left and right images; see Fig.S8 and Fig.S9 for more seeds). In this case, no continuity is imposed on the hidden layer, where the weights can either be factorized only or free-kernels. By factorized only we mean that the rotations array is not encoded by an R-DPPN but is trained directly. For factorization, a random non-differentiable translation (an additional

affine transformation) was applied to the weights during training to enhance its degrees of freedom compared to the free-kernels case. By comparing the two maps (Fig. 3c left and right images), it can be seen that factorization, aside from being more efficient, leads to cleaner map formation than free-weights. This can be inferred from the more precise delineation of the orientation domains in the factorization case. One possible reason for this difference is that the factorized weights/filters seem to have narrower or more monotonic tuning curves than the free-weights case as seen by comparing the weights in each case, and by inspecting the tuning curves (Fig. 3d; see Fig.S10 for more tuning curves). From these results, we can see that the tuning curves of the model with free-weights can be wider and/or multimodal.

To summarize the tuning curves, we calculated the distribution of their widths (Fig. 3e with equation (E1)). From these results, we can see that the tuning curves of free-weights are wider than that of factorization. The left most red bar in the free-weights case is due to zero activations (inactive) neurons in the hidden layer. This is due to the weights of these neurons being negative while the activation function is Relu (for example, Fig. 3c right image, middle left kernel is all negative).

#### 3.1.4 Spatial frequency selectivity is inversely proportional to the receptive field size

The human visual system is scale invariant, namely, it can recognize patterns across different spatial scales (Han et al., 2020). It is hypothesized that this scale invariance can be supported by the receptive field properties in V1. Specifically, different neurons respond to different spatial frequency bands. In addition, it was found that this frequency tuning is inversely proportional to the receptive field size (Teichert et al., 2007; Wiecek et al., 2026).

Here, we show that this relationship between receptive field size and frequency tuning can arise in networks that utilize factorization in the hidden layer (Fig. 3f). Specifically, during training we applied two non-differentiable affine transformations, scaling and translation, to the weights. The resulting filters showed orientation and spatial frequency tuning. In addition, the frequency tuning was inversely proportional to the receptive field size (Fig. 3f right images).

### 3.2 Generalization

Recognizing the properties and affordances (classification) of an unfamiliar object is a form of generalization that humans and some animals possess. Here, we show that this kind of generalization is possible through the combination of a factorial code and spatial encoding. A factorial code decomposes the object into a set of basic features, while spatial encoding learns a smooth map in the output features that allows for the recognition of unfamiliar features, and subsequently novel feature combinations. It is hypothesised that this factorial code might be utilized by the hippocampus and other areas like the prefrontal cortex (Bernardi et al., 2020; Villalba et al., 2025). Thus, as another example of how our setup is favourable for encoding innate functions, we study the problem of postnatal generalizations that are based on a factorial code.

We demonstrate how the genomic code, with the aid of factorization and spatial encodings, can efficiently initialize brain networks that can process novel postnatal feature combinations. This is achieved by utilizing a range of network architectures (Fig. 4a) and tasks (Fig. 4b). The network has one hidden layer, where the hidden weights are regularized by factorization (FT) + R-DPPN or being trained freely (FW). The final layer is made up of two DPPNs; one for each feature. Thus, if more features are being learned, more DPPNs are needed. The problem set is composed of three problem types, where in each problem, two features are varied. These problems are i) *Circle*, where we vary the scale of the circle and the orientation of the opening. ii) *Arc*, where we nonlinearly scale the object, while also varying its orientation. iii) *Comb*, where we vary the number of teeth in the object while also varying its orientation (see section 2 and section E1 for more details).

**Fig. 4:**
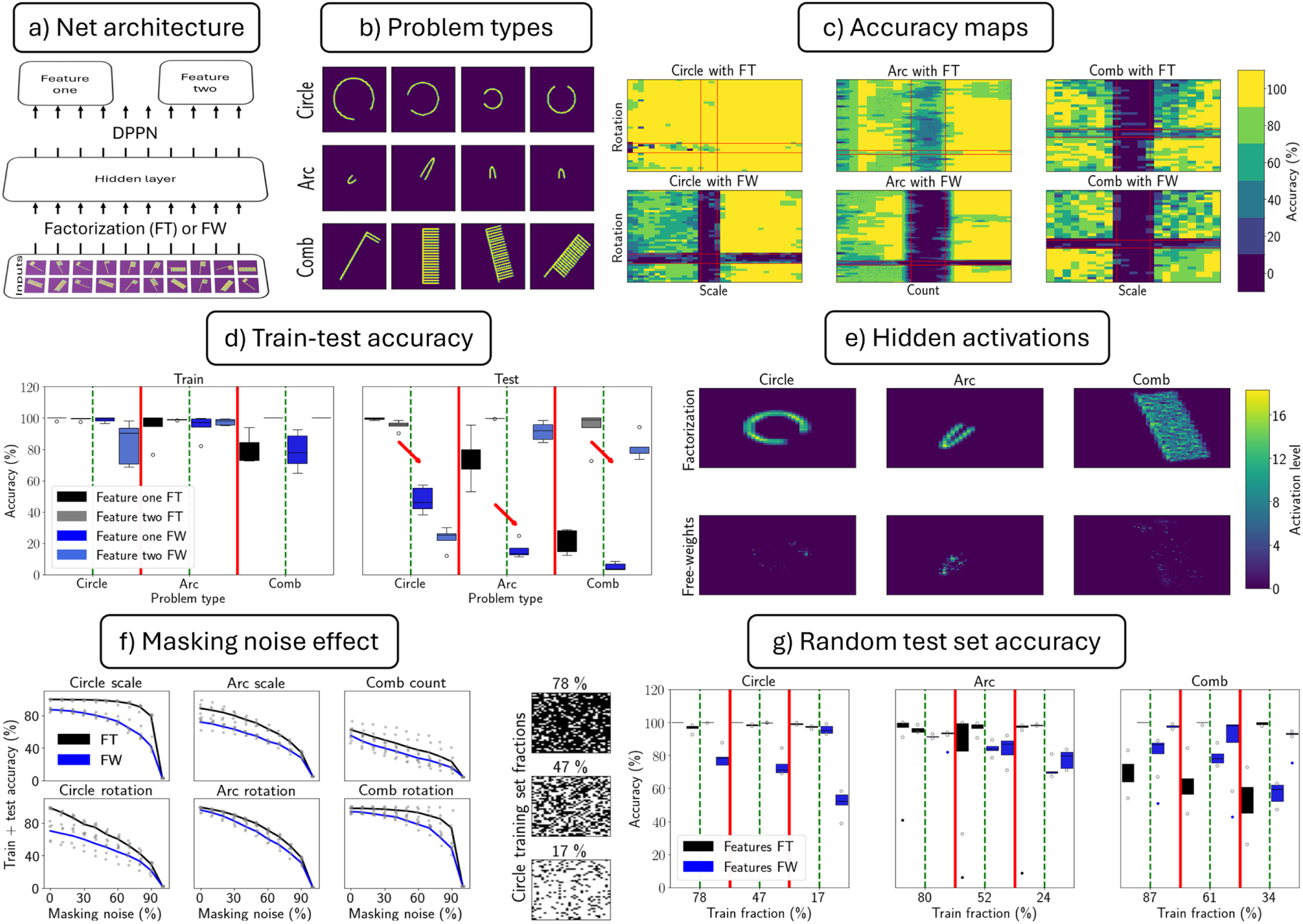
Genomic feature generalization. **a)** Network Architecture: the hidden layer can be factorized-weights (FT) + R-DPPN or free-weights (FW), while the final layer is made up of two DPPNs. **b)** Problem Types. We employ three types, *Circle*, *Arc* and *Comb*. For each type, we vary two features, scale and rotation for Circle, non-linear scaling and rotation for arc, and tooth count and rotation for Comb. **c)** Accuracy Maps: each point is the accuracy over 5 seeds for a particular output class (feature combination). **d)** Summary of train and test accuracy for each problem type and hidden layer architecture. The red lines separate the problem types, while the green lines separate the hidden layer architectures. **e)** Hidden activations for each problem type and hidden layer architecture. **f)** Masking noise effect: inputs were multiplied with a mask of 0s and 1s sampled from a uniform distribution with the 0 probability presented on the *x*-axis. **g)** Sparse training sets accuracy: training examples are randomly chosen from the input data set with varying degrees of sparsity rather than using pre-specified, contiguous inputs. The red lines separate the various training set fractions, while the green lines separate the hidden layer architectures. The lone dots are outlier data points.

#### 3.2.1 DPPNs can feature generalize with the appropriate output code

Using a factorial code in the output layer, we assess the generalization ability of our network by testing on unseen feature combinations during training (Fig. 4c regions demarcated by red lines). From these tests, we observe two things: i) the network can generalize to unseen combinations of features (Fig. 4c top row), and this can be attributed to the DPPN’s ability to learn smooth weight structures (see Fig.S13). ii) the generalization ability is dependent on the problem type, where in terms of success, *Circle > Arc > Comb*.

#### 3.2.2 Weight factorization is efficient and favourable for feature generalization

When comparing the effect of the hidden layer architecture on performance (Fig. 4c), we observe that factorization is more advantageous to generalization than free-weights for all problem types. This discovery is also evident in the train-test accuracy summary (Fig. 4d), where we see that the

test accuracy goes down, marked by the red arrows, as we switch from factorization to free-weights in the hidden layer. The success of factorization over free-weights can be attributed to the nature of the hidden layer activations (Fig. 4e), which are a result of the learned weights. From these activations, we see that factorization learns equivariant representations, where the general features of the object are preserved under transformations. While for the free-weights case, the network learns some kind of unique instance-based representation. Regarding efficiency, the hidden layer with factorization uses only ≈ 150 parameters (≈ 50 for the kernels and 100 for the R-DPPN), while the free-weights case uses in the order of 10^5^ parameters.

#### 3.2.3 Weight factorization is more robust to masking noise

Another advantage of factorization over free-weights is its robustness to masking noise as evident from inspection of the train+test datasets (full dataset) accuracy (Fig. 4e). Masking noise is a form of multiplicative noise where the input to the network is multiplied with a mask of 1s and 0s with varying probability of 0s. Weight factorization shows more robustness to this form of noise in two ways: i) at each noise level, factorization has higher accuracy than free-weights for all problem types; ii) in some cases, factorization is resilient to gradual increases in noise levels as seen from the top left and bottom middle plots (Circle scale and Comb rotation). In these plots, as the noise level increases, the accuracy gap can be seen to increase until both drop to chance level at very high noise levels.

#### 3.2.4 Genomic generalization is robust against a sparse distribution of training data

A more natural train/test division is when the training set is chosen at random and sparse. Thus, we tested each network’s ability to generalize under such conditions and found that the networks can be resilient to a degradation in accuracy, even at very sparse training sets (see Fig. 4g, where example training set fractions from the Circle case are shown on the left). This resilience might be attributed to the fact that individual features (not feature combinations) are observed more than once. In addition, factorization still outperforms free-weights in tests of generalization, except for two cases (Fig. 4g most right plots, 87% and 61% in the Comb problem).

### 3.3 Efficient genomic encoding

In this section, we asked if there is a selection pressure on the size of the genome. Experimentally, it has been shown that there is correlation between the genome size and the ability to reproduce in seed beetles (Arnqvist et al., 2015). Also, it was demonstrated that in microalgae there is a conflict of interest regarding the genome size: smaller sizes are needed to improve performance and minimize unnecessary DNA, while the genome still needs to be large enough to support important functions (Malerba et al., 2020). Although these studies are limited to certain organisms, a hypothesis can still be formulated about the requirements for the genome’s size: the genome needs to efficient, in the sense that it makes good use of every base pair. For example, hot peppers have on the order of ≈ 3.5 *billion* base pairs (Hulse-Kemp et al., 2018), while humans only have ≈ 3 *billion*, and it can be argued that humans are more intelligent than hot peppers. Thus, it can be postulated that a requirement of the genomic code is the efficient compression of information which we investigate here.

We show that one of the highest forms of compression of a genomic code can be acquired by spatial encodings of factorizable parameters, like weights. We support this claim with the aid of the MNIST Benchmark and a collection of diverse network architectures (Fig. 5a). In these networks, we vary the hidden layer count and the kernel size in each layer, while the final layer is a DPPN with varying hidden sizes. However, the kernel sizes are fixed for the same network, i.e., no change in kernel size with depth. Each hidden layer is made of factorized weights. Namely, a hidden layer is made up of a canonical kernel and an R-DPPN that specifies its rotations. For the convolution operation with input and activations, we use a valid padding to counter the effect of the increase in hidden size due to afferent expansion (see section 2 and section E1 for more details).

**Fig. 5:**
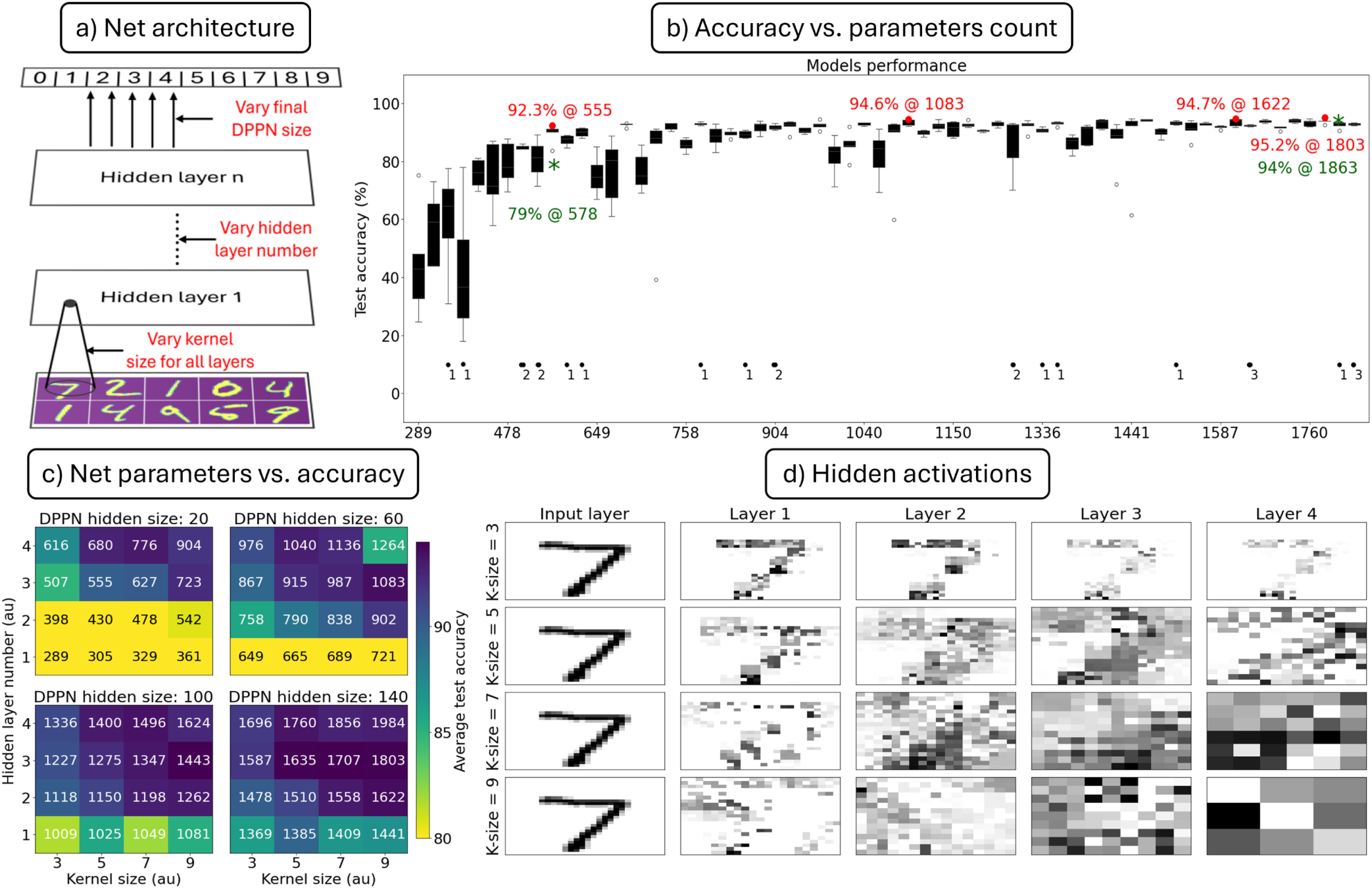
Genomic efficient coding. **a)** Network Architectures: we use weight factorization in the hidden layers. The hidden layer number varies between 1 and 4 with kernel sizes ∈ {3, 5, 7, 9}, while the final layer is a DPPN with one hidden layer, but of varying size. **b)** Accuracy as a function of networks parameter count. Chance accuracy, where a network fails to learn, is shown at the bottom with the associated trial count out of 5 seeds. Some of the best performing networks are highlighted in red with some results from literature in green asterisks and text. **c)** Average test accuracy as a function of the DPPN hidden size, number of hidden layers and kernel sizes. The parameter count is embedded in each cell. **d)** Hidden activations as a function of kernel size for networks of 4 layers. Each activation image has its own scale, as the emphasis here is on the relative strength of the activations i.e. the activations structure.

#### 3.3.1 Factorization plus spatial encodings might offer the highest form of genomic compression

By combining factorization with DPPNs, we can reach near peak performance (95.2%) with only ≈ 1, 800 parameters (Fig. 5b). From studying the relationship between performance and parameter

count, we find that the test accuracy reaches a cap at around 95%, and it can be argued that this accuracy cap is a reasonable limit given how deformed some of the MNIST digits are.

For comparison with other findings, we also highlighted some of the best model test accuracies in red and contrasted them with some results from the literature in green (Fig.5b). Specifically, we added two results from (Shuvaeva et al., 2024) on MNIST: i) 94% with 1, 863 parameters (322 fold from 6 × 10^5^) and ii) 79% with 578 parameters (1, 038 fold from 6 × 10^5^). These two cases are highlighted as green text and asterisks on the very top right and on the left respectively. In addition, for the first layer in the phenotype, some of the canonical filters used in factorization resemble line orientation filters (Fig.S14).

#### 3.3.2 Layer number and DPPN hidden size have more impact on performance than kernel size

Increasing the final DPPN hidden layer size increases the test accuracy across various configurations (Fig.5c). This outcome might be due to a decrease in underfitting, as larger encoding networks are needed for better input-output mapping in the final layer. In addition to the influence of the final DPPN, the number of hidden layers in the phenotype also has a significant impact on performance which might be attributable to two things: i) better feature extraction, and ii) feature map compression. The evidence for feature extraction can be supported by the near preservation of the hidden representation dimensions while the hidden activations structure changes across layers (Fig.5d top row). Feature extraction is a possible reason for why final DPPNs with small hidden sizes have higher performance in phenotypes with 3+ layers (Fig.5c top left image). In the context of spatial encodings, feature compression is beneficial since it reduces the number of weights needed in the final layer, and subsequently leads to smaller encoding networks. However, too much compression might be disadvantageous as retinotopy and distinctions between classes are smeared (Fig.5e bottom right images).

## 4 Summary and discussion

We propose a novel hypothesis explaining how the genomic code, despite its limited informational capacity, efficiently initializes innate brain functions comprising billions of neurons through two complementary mechanisms: spatial encodings and factorization. Spatial encodings, in the form of differentiable pattern-producing networks (*DPPNs*), are network types that indirectly encode the weights of another network (referred to as a hypernetworks in machine learning). Using this formalism, the spatial encoding networks act as the genotype and the encoded networks as the phenotype. Spatial encodings serve two primary functions: i) they compress the parameter size of the phenotype as they themselves can be of much smaller size and ii) they enforce continuity on the weight structure because they encode the weights as a function of the associated positions of the pre- and post-synaptic neurons, such that nearby neurons have nearby weight values.

Factorization, as introduced in this work, is a method by which we decompose the weights of a layer into its constituent interacting ingredients. Specifically, we abstract (factorize) the layer’s weights into a spatially independent weight kernel (*canonical filter*) and its spatially dependent affine transformations along the spatial dimensions of the layer. In this work, we limit the differentiable transformations to only rotations. Factorization and DPPNs can interact by using DPPNs to encode the rotations of the canonical filter, thus enforcing continuity on the rotations while compressing their representations. Thus, we can use DPPNs in two ways, either to encode the weights of a layer, or to encode the rotations of a factorized layer. In the case of encoding rotations, we name them *R-DPPNs*. Using these mechanisms of factorization and spatial encodings, we have provided an account of how the genomic code can efficiently initialize innate functions such as V1 orientation maps and networks with the ability to generalize.

For V1 maps, we have shown several novel ways by which these maps can emerge: i) by using a DPPN to encode the hidden layer, ii) by using factorization + R-DPPN to encode the hidden layer, iii) by using a DPPN to encode the final layer, and iv) by using factorization + a Gaussian kernel in the hidden layer. The first three use indirect encoding, while the last uses direct encoding. For all cases, we also get weight kernels that resemble line filters. One possible reason for the emergence of such maps and filters is the interaction between the need to represent every possible feature (line orientation) at each spatial position while also enforcing continuity in the activations profile by the DPPNs, R-DPPNs and Gaussian kernels. In addition, the applied continuity also affects the final layer’s weights as seen from the structural correlations between the emergent maps and these weights (Fig.S11).

To assess the benefits of using factorization in the hidden layer, we compared two cases: i) the hidden layer is encoded by a DPPN, or ii) it is encoded by factorization + R-DPPN. Both indirect encoding networks — DPPN and R-DPPN — had one hidden layer with a varying number of neurons. For all hidden sizes (except the case of one neuron), we found that factorization + R-DPPN, on average, are more efficient in their encoding as they use fewer parameters, they have slightly higher performance, they converge faster (Fig.S4, Table S1), and their encoded weight structures have higher directionality. However, the only drawback to factorization is that the emergent maps do not exhibit the full range of biological features. Although we observe continuity and iso-orientation domains, we do not find the characteristic pin-wheels. This structural limitation likely stems from the R-DPPN mapping spatial positions directly to continuous scalar rotations, creating a framework where converging to a complex pin-wheel topology is structurally penalized when simpler, efficient alternative solutions exist.

In general, the proposed approach for explaining the emergence of orientation maps is unique in two key aspects. First, it is more biologically plausible in that we did not use an explicit spatial correlation loss function to enforce continuity in a backpropagation framework (Shuvaeva et al., 2024). However, a recent finding (Qian et al., 2026) showed that V1-like orientation domains can arise in a backpropagation framework when incorporating lateral connections. Despite this, our approach stands out in that it provides a developmental account through indirect encoding and is more efficient. Secondly, it might be the first case where maps emerged without employing isotropic continuity, like isotropic Gaussian functions in a learning framework, to motivate the emergence of maps, which is the traditional approach in self-organizing models. Furthermore, our framework represents, to our knowledge, the first instance of orientation maps concurrently emerging alongside edge and line filters within a backpropagation regime.

Furthermore, for V1 maps, we showed that when we applied non-differentiable affine translation and scaling, we replicated the inverse relationship between the receptive field size and spatial frequency tuning. This was achieved with the factorization framework, namely with a canonical filter that is differentiably rotated along the spatial dimensions of the layer. Consequently, different V1 neuronal responses can be attributed to the same mechanism (a single canonical filter) but with different receptive field sizes. This kind of minimal coding might be beneficial for an evolving genome with limited informational capacity.

Beyond explaining the emergence of V1 maps, our approach demonstrates how innate functions with the ability to generalize can be encoded. This was illustrated by utilizing datasets that have dual geometric features. These datasets were comprised of Circles with varying scale and orientation, Arcs with varying non-linear scale and orientation, and Combs with varying tooth count and orientation. Utilizing these datasets, we showed that after training, a phenotypic network can generalize to unseen feature combinations by applying a DPPN to the final layer. Furthermore, we found that this ability to generalize is dependent on the problem in question. This dependence on problem type might be attributed to the nature of intra-dataset transformations, and these transformations can be divided into two subtypes, linear and non-linear. Rotation is linear, scaling is linear, while tooth number is not. However, the non-strictly linear scaling of the Arc problem may account for its slight performance degradation compared to the Circle problem.

Beyond the necessity of having a DPPN in the final layer to generalize, we also showed that factorization is advantageous and more efficient than free-weights in the hidden layer. This advantage stems from the emergent activations; namely, factorization leads to equivariant representations that support generalization, while free-weights do not. In addition, we showed that factorization is more robust and resilient to masking noise for all problem types and features. However, when tested with a random test set, factorization showed an advantage over free-weights for two out of the three datasets, namely Circle and Arc.

From these results on generalization, it can be concluded that under the factorial hypothesis, where objects are recognized by their features, the genome employs techniques that restrict and regularise the change in learned features across the cortical spatial dimensions in a smooth and predictable manner that enhances downstream higher-level functional learning. In other words, the genome learns a smooth map in feature transitions to support generalization in higher-level functions, and the emergence of such maps can be enhanced with the utilization of factorization and spatial encodings. From a modelling perspective, these results suggest that relevant structure and biases are needed in neural networks to simulate some aspects of human cognition. This can be contrasted with conventional ANNs that fail on object recognition when confronted with simple image manipulations (Su et al., 2019).

Finally, we have shown that applying factorization + R-DPPN in the hidden layer, while having a DPPN in the final layer might be one of the highest forms of genomic compression for arbitrary innate functions. This was exemplified by the MNIST dataset. Though we have shown that we have achieved higher performance than recent results (Shuvaeva et al., 2024), our accuracy hits a ceiling around ≈ 95%. This seemingly hard limit on the maximum performance might be attributed to the difficulty of classifying some digits that are highly deformed.

From all of the above findings, we can make more detailed claims on how the genomic code can benefit from both spatial encodings and factorization. Spatial encodings, in the form of a DPPN or an R-DPPN, contribute two things, compression and continuity. Regarding compression, the efficiency of the compression depends on the regularities of the encoded structure, and by structure we mean a layer’s weight matrix or an array of rotations. Highly regular structures can be compressed more efficiently than irregular ones (Clune et al., 2011). Thus, if a structure is highly irregular, a larger DPPN is needed to capture its irregular structure. However, this relationship between DPPN size and complexity can be decoupled by applying factorization. Factorization adapts and enforces a weight structure that is independent of how complex the underlying input structure is. Namely, the weights will always be a canonical filter that is spatially rotated irrespective of complexity of the input. Subsequently, this will reduce the complexity of the problem to be encoded by a DPPN. Before factorization, a DPPN had to encode the full underlying structure, but with factorization it need only encode a 2D array of rotations as an R-DPPN. This simplification of the problem has a cost in the form of reduced performance regardless of the size of the R-DPPN. However, from an evolutionary perspective, performance is not the target of the genomic initialization, the goal is providing an acceptable initial state for in-life learning. Finally, factorization also has the advantage of reducing the variance in the weight structure, which is beneficial when equivariant representations are desired.

### 4.1 Limitations and future work

In this work, we utilized simple artificial input datasets to demonstrate, from first principles, the underlying mechanisms and dynamics of our approach. Thus, a natural next step is to utilize more realistic input datasets, like natural images. In the context of V1 maps, if natural stimuli are used, we can investigate the nature of the weight structure and computational maps that arise in a stack of multiple hidden layers as may be required by more complex inputs.

For our factorization approach, it is very restrictive in that it only considers rotations, and this can be expanded on in several ways. For example, adding parallel factorization layers. In this context, the question is: are parallel maps of affine transformations sufficient to fully learn a problem? Another direction of expansion is applying non-linear transformations instead of the affine (linear) ones. Also, regarding the comparison between factorization + R-DPPN and DPPN, more comprehensive studies are needed in terms of more input problems with diverse structures. In addition, it would be more naturalistic to compare both of these approaches in an evolutionary framework.

Finally, the factorial code utilized in this work is manually constructed to be compatible with the corresponding dataset. Thus, an intuitive next step is to find a way to learn the factorial code in an unsupervised way. In addition, for the factorial code itself, it would prove useful to more extensively study the relationship between the nature of the feature transformations and its ability to be generalized.

## Author contributions

Jeffrey S. Bowers, Benjamin D. Evans and Dan F. M. Goodman supervised the work, analysed the data and revised the manuscript. Karim G. Habashy conducted the research and drafted the manuscript.

## E1 Extended Methods

For all differentiable parameters in this work, we used different learning rates for direct weights vs. DPPNs. Canonical filters, arrays of transformations or filters, used a learning rate of 0.01. While, any indirect encoding networks, like DPPNs and R-DPPNs, used a learning rate of 0.001. These learning rates were used with the optimizer AdamW in the PyTorch Python library. The only exception to this is when we compared DPPN to factorization + R-DPPN. In this case, we used a 0.001 learning rate.

### E1.1 Problem sets

Here we present more details about the problem sets used in this work.

#### E1.1.1 Efficient coding

The problem set is the standard MNIST dataset with 6 × 10^4^ and 1 × 10^4^ train-test division. The batch size was 1000 for training, while testing was done with the full test size.

#### E1.1.2 Maps

We used as input lines with varying orientations, positions, lengths and thicknesses. The orientation angle step was 20*^o^* spanning the range 0*^o^* − 180*^o^*. The thickness ∈ {1, 2, 3}, while the length ∈ [5, 15] with a step of 1. For Fig.3f, to test the orientation and frequency tunings of the receptive fields, we applied sinusoidal gratings to the receptive fields with spatial frequencies ∈ [2, 8.8] with a step of 0.2, and angles [0, 160] with a step of 20.

#### E1.1.3 Generalization

The following applies to all figures except Fig.4g, where we used a randomized training set. For all problem types, we used an input size of (121, 121) and line thickness of 2. For the Circle problem, we had 32 continuous variations in scale and 41 continuous variations in rotation. The test set was acquired by removing features in the index range (13, 14, 15, 16) from scale and (8, 9, 10, 11, 12) from rotations, giving us a test fraction of 23% For the Arc problem, we had 20 continuous variations in scale and 72 continuous variations in rotation. The test set was acquired by removing features in the index range (8, 9, 10, 11, 12) from scale and (13, 14, 15, 16) from rotations, giving us a test fraction of 29%. For the Comb problem, we had 21 continuous variations in pin count and 36 continuous variations in rotation. The test set was acquired by removing features in the index range (8, 9, 10, 11, 12) from pin count and (13, 14, 15, 16) from rotations, giving us a test fraction of 32%.

### E1.2 Extra equations

#### E1.2.1 Tuning width calculation

To measure the width of a tuning curve, we integrated the firing rates weighted by their distance from the maximum using (E1)

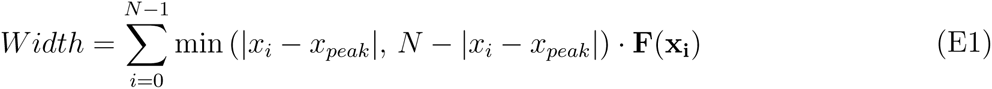

In this equation, *x_peak_* is the index (location) of the maximum firing rate. *x_i_* is the index of a particular firing rate. **F**(**x_i_**) is the firing rate at location *x_i_*. *N* is the size of the tuning curve (number of orientation angles).

#### E1.2.2 Parameter count calculation

To quantify the parameter efficiency of a model we count the number of parameters in a network using equation (E2)

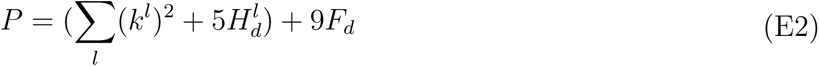

In this equation *P* is the total parameter count in the network. *k^l^* is the kernel size in layer *l*. *H_d_*is the phenotypic hidden layer R-DPPN hidden size. *F_d_*is the phenotypic final layer DPPN hidden size. *H_d_* is always fixed at 20. Thus, for a 1 layer network with 3 × 3 kernel and final DPPN hidden size of 20, we have a parameter count of 289, which is the smallest network we employ as seen from the start of the *x*-axis in Fig.5b.

### E1.3 Network parameters

Here we summarize and show more details about the various network parameters for all networks utilized in this work. This is shown in Table E1.

## S1 Supplementary Material

### S1.1 V1 orientation maps

In this part of the supplementary material, we show some data referred to in the main text. First we show different seeds for the DPPN result in Fig. 2b in the main text (Fig.S1). Next, we show other seeds for the Factorization + R-DPPN case in Fig. 2b in the main text (Fig.S2).

We also show a more detailed comparison between DPPN and Factorization (Fig. 2b main text, left and right images) for the generation of V1 orientation maps as a function of the hidden layer size (Fig.S3 and Fig.S5). This comparison includes parameter count, performance, weight structure, directionality and convergence.

Parameter count is the number of parameters used in the hidden layer. Performance is the average loss in the final 2000 training epochs where learning is frozen in this interval. Weight structure involves the shape and weight values distribution. Directionality is a measure of the sharpness of the orientation of the weight kernel. For example, a circular Gaussian blob would give a low directionality, while an ellipsoidal shape with large axis ratio (long/small) would give high directionality. Convergence is how fast a particular loss value is achieved during training (Fig.S4). A summary of how these parameters are influenced by the hidden architecture and hidden size is provided (Table S1). For the Factorization+DPPN case, there are two outliers in the loss data for the hidden sizes 2 and 3 respectively. These outliers are included in the mean loss calculations, but omitted from the convergence calculations.

Another example of how factorization and continuity can lead to the emergence of V1 orientation maps is introduced here. We show how coupling factorization and lateral Gaussian kernel can lead to the emergence of V1 orientation maps (Fig.S6). The factorization in this case has no R-DPPN, but continuity is enforced by the Gaussian kernel. The Gaussian kernel is of size 11 and is applied iteratively three times on the hidden activations, then the output is passed on to the final layer. The kernel itself is a mixture of Gaussians of opposite signs, but with different spatial coefficient (or standard deviations), these are 0.2 and 2.4 respectively. The resultant kernel is scaled by a scale of 20. We also show other seeds of this configuration (Fig.S7).

For Fig. 2c in the main text, were we have a fully connected layer as the final layer, we show other seeds for the Factorization case (Fig.S8), and the Free-weights case (Fig.S9).

We also show other tuning curves for the neurons/maps shown in Fig. 2c of the main text (Fig.S10).

Regarding the map structure in the hidden layer, it is observed that the map structure correlates with the final layer weights (Fig.S11). All nine output neurons have weights that resemble the hidden layer activations.

Finally, we show that larger kernel sizes give rise to more complex receptive field structures (Fig.S12).

### S1.2 Generalization

In this part of the supplementary material, we show the final weights structures learned by the DPPN in Fig. 3c of the main text (Fig.S13).

### S1.3 Efficient genomic encoding

In this part of the supplementary material, we show the properties and shapes of the learned weights in the “efficient genomic encoding” section. Specifically, We show all the learned canonical filters across all trials for layer one (Fig.S14). Canonical filters are filters pre-transformations and they are fixed for the whole layer. One gets a whole layer by rotating these filters at various spatial locations. It can be seen that a majority of the canonical filters take the form of an edge or line filter. However, for the second layer and above, it is hard to discern the utility of the learned filter, and thus they were not shown.

---

**Fig. S1:**
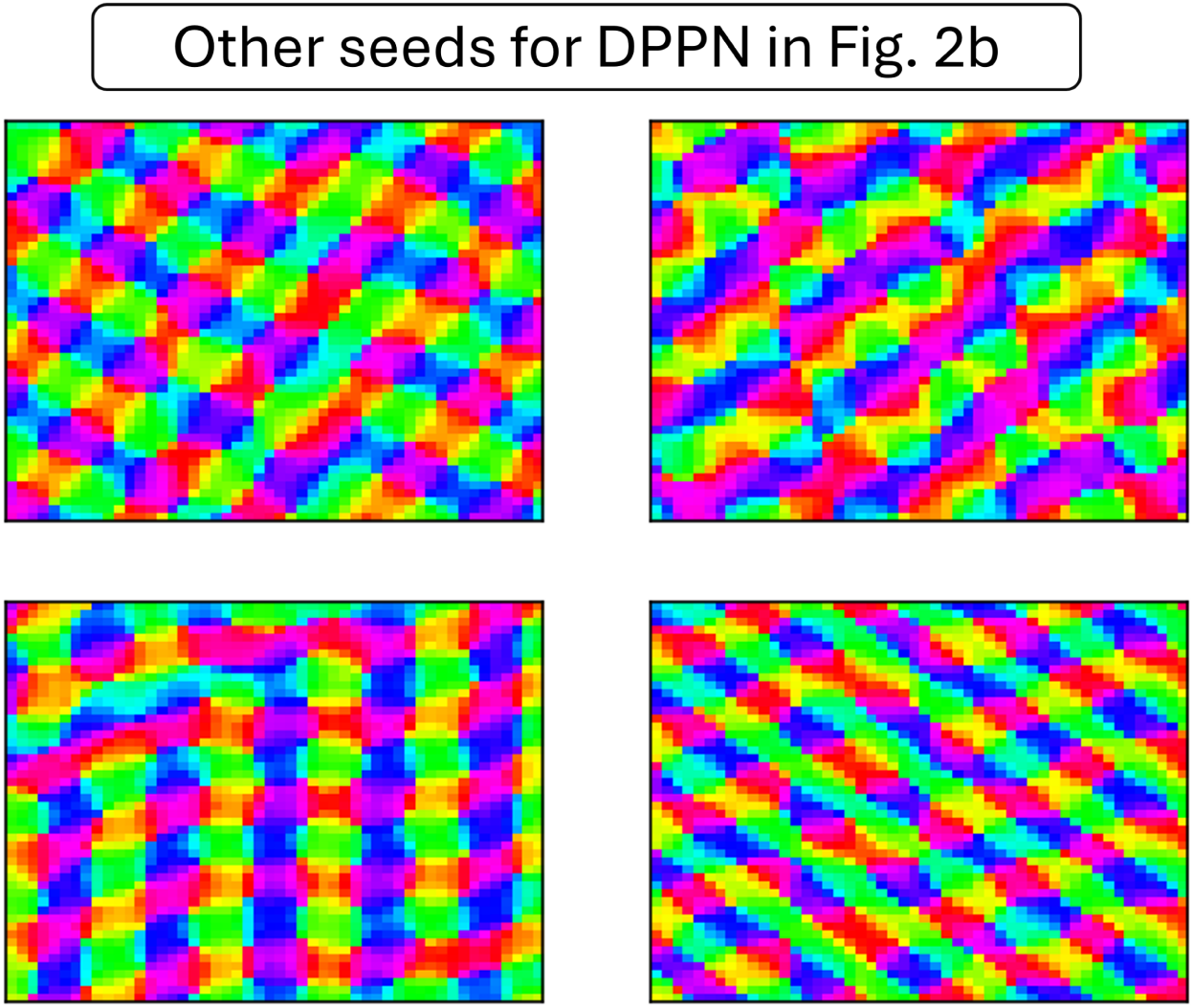
Other seeds for the DPPN case in Fig. 2b in the main text.

**Fig. S2:**
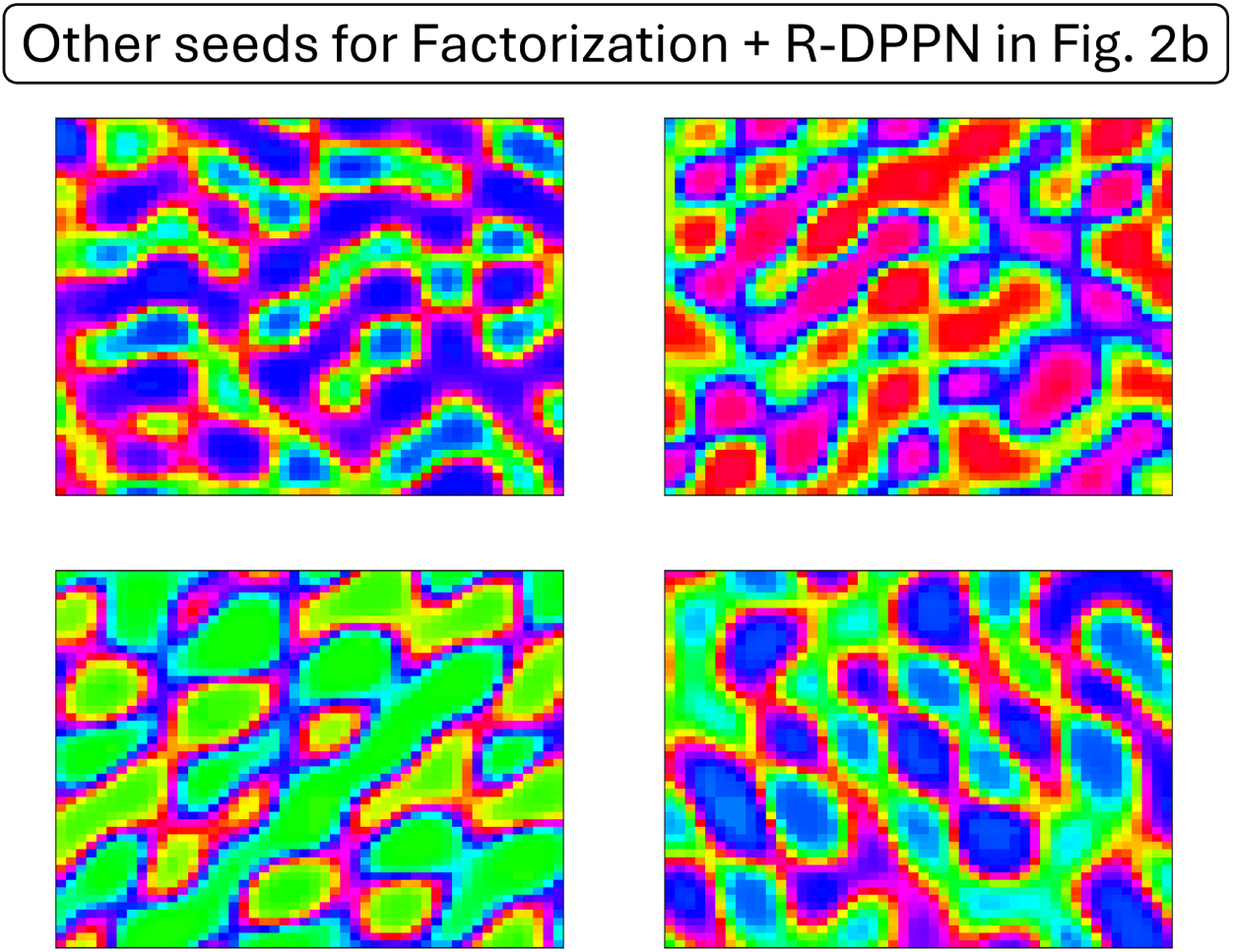
Other seeds for the Factorization + R-DPPN case in Fig. 2b in the main text.

**Fig. S3:**
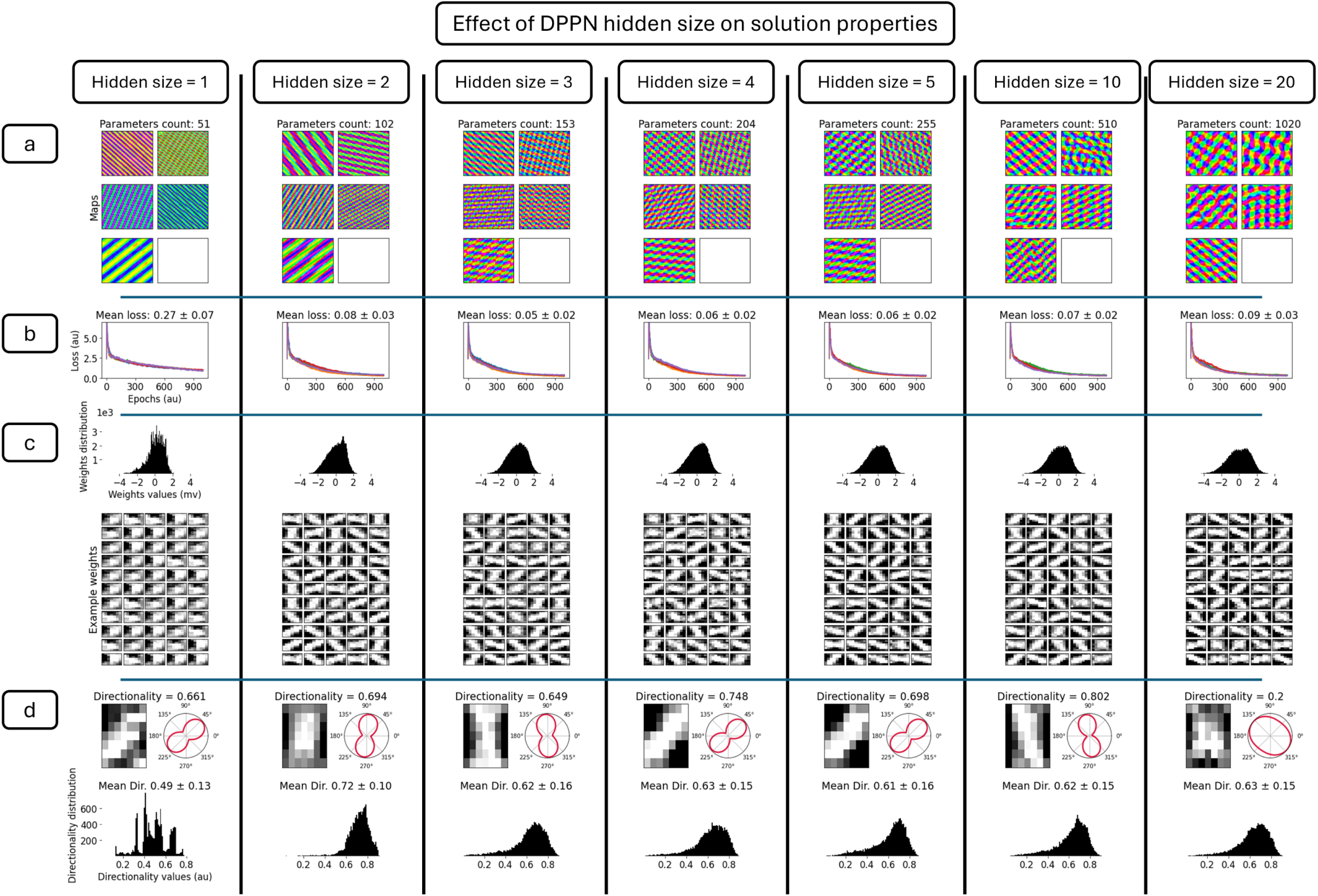
More details on the maps generated by the DPPN case (Fig. 2b left) for various DPPN hidden layer sizes. **a)** Maps as a function of the hidden size. Here we show 5 seeds for each map. **b)** training loss for all seeds. **c)** weight values distribution for all seeds. We also show example weight kernels. **d)** directionality, where in the top row we show example weight kernel and its directionality value. In the bottom row, we show the directionality distribution for all seeds.

**Fig. S4:**
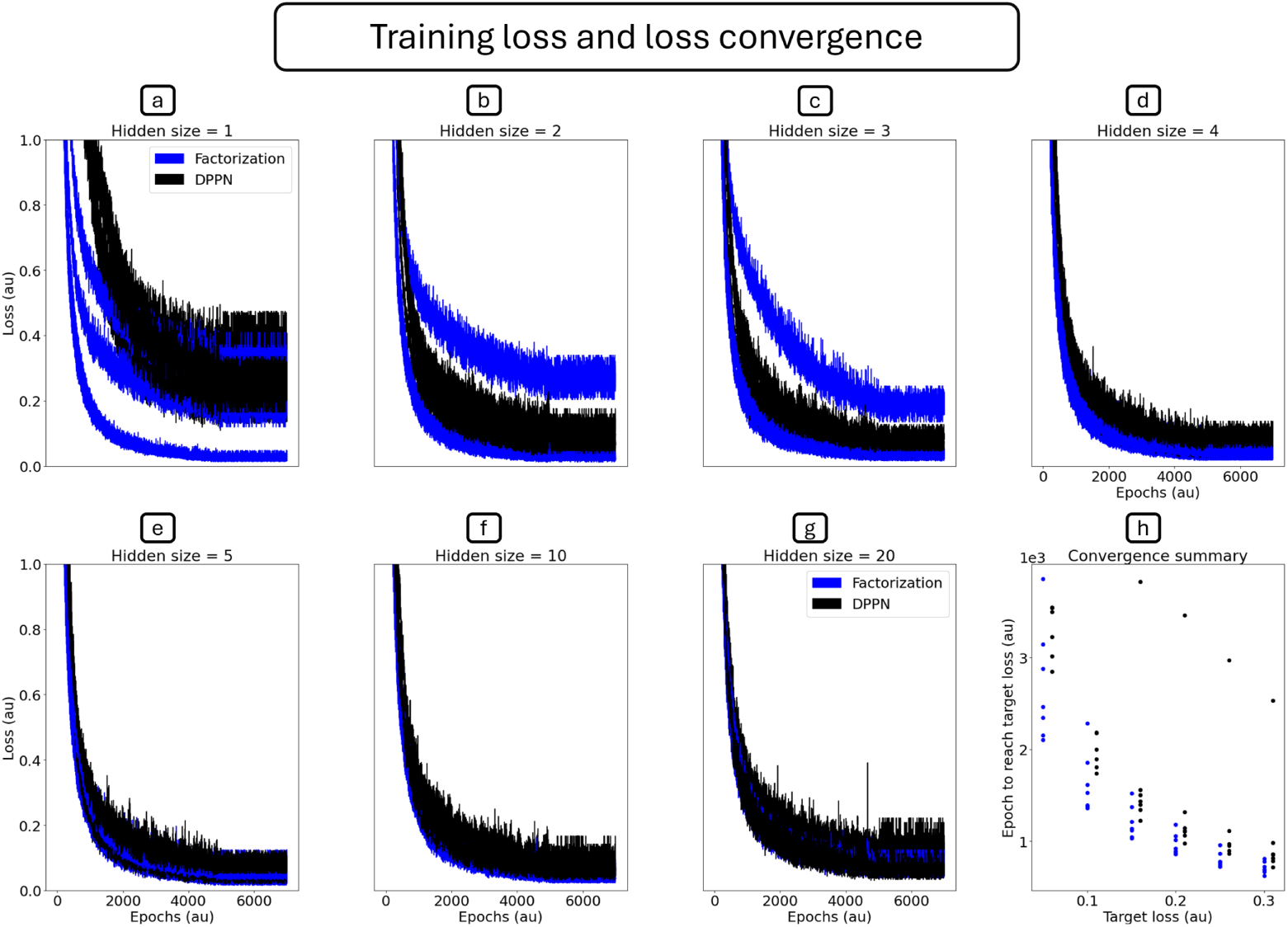
Training loss and loss convergence. **a)** to **g)**, training loss as a function of hidden size. **h)** Loss convergence summary, where we extract the number of epochs needed to reach a target loss.

**Fig. S5:**
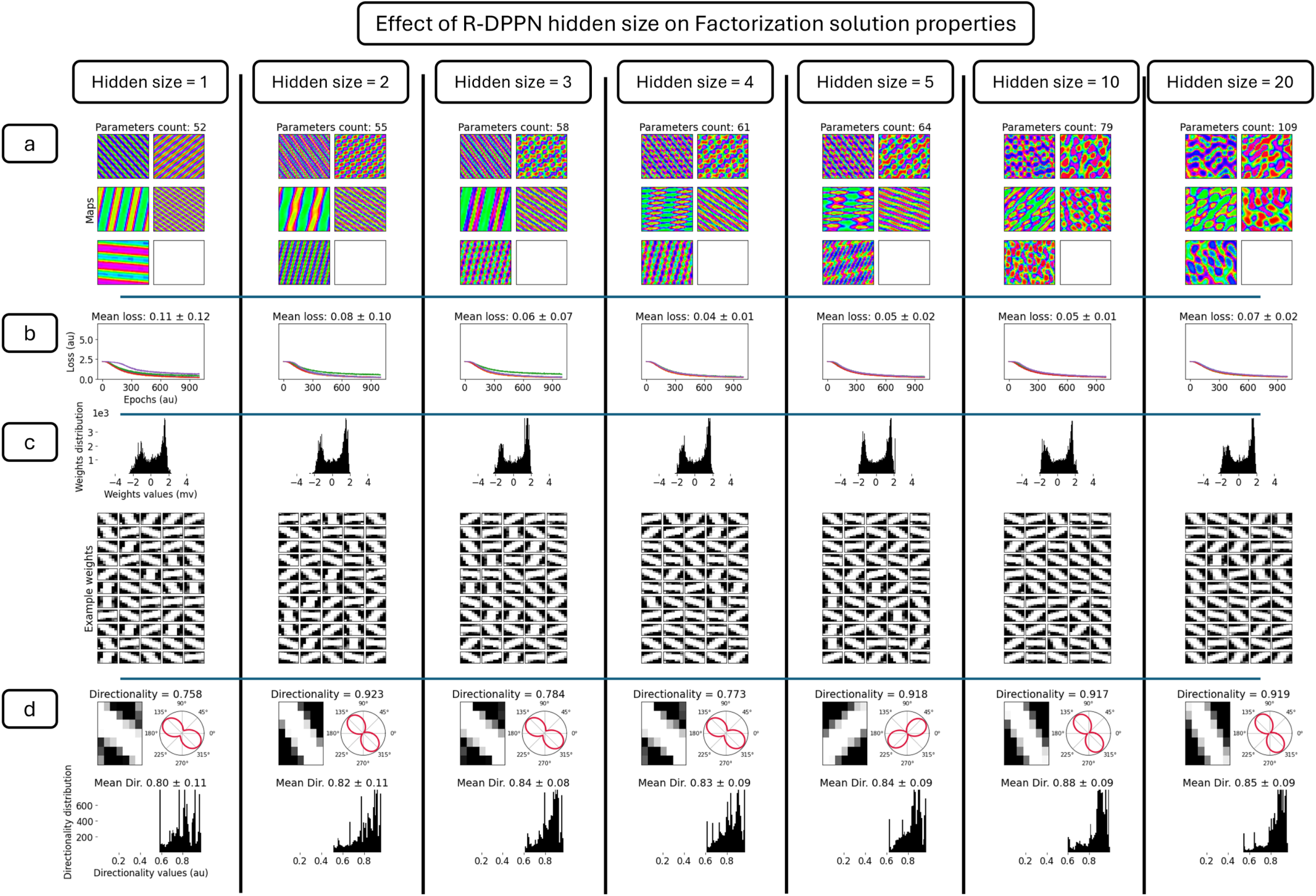
More details on the maps generated by the Factorization + R-DPPN case (Fig. 2b right) for various R-DPPN hidden layer sizes. **a)** Maps as a function of the hidden size. Here we show 5 seeds for each map. **b)** training loss for all seeds. **c)** weight values distribution for all seeds. We also show example weight kernels. **d)** directionality, where in the top row we show example weight kernel and its directionality value. In the bottom row, we show the directionality distribution for all seeds.

**Fig. S6:**
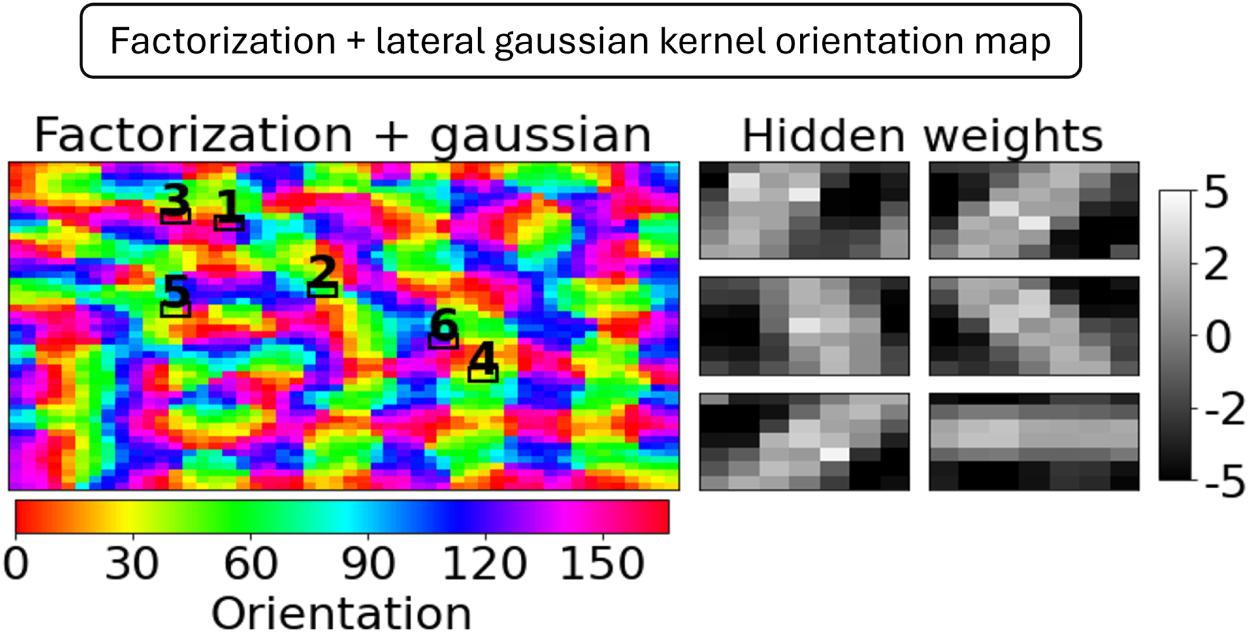
**V1 orientation maps emerge when combining factorization and a lateral Gaussian kernel that enforces continuity.**

**Fig. S7:**
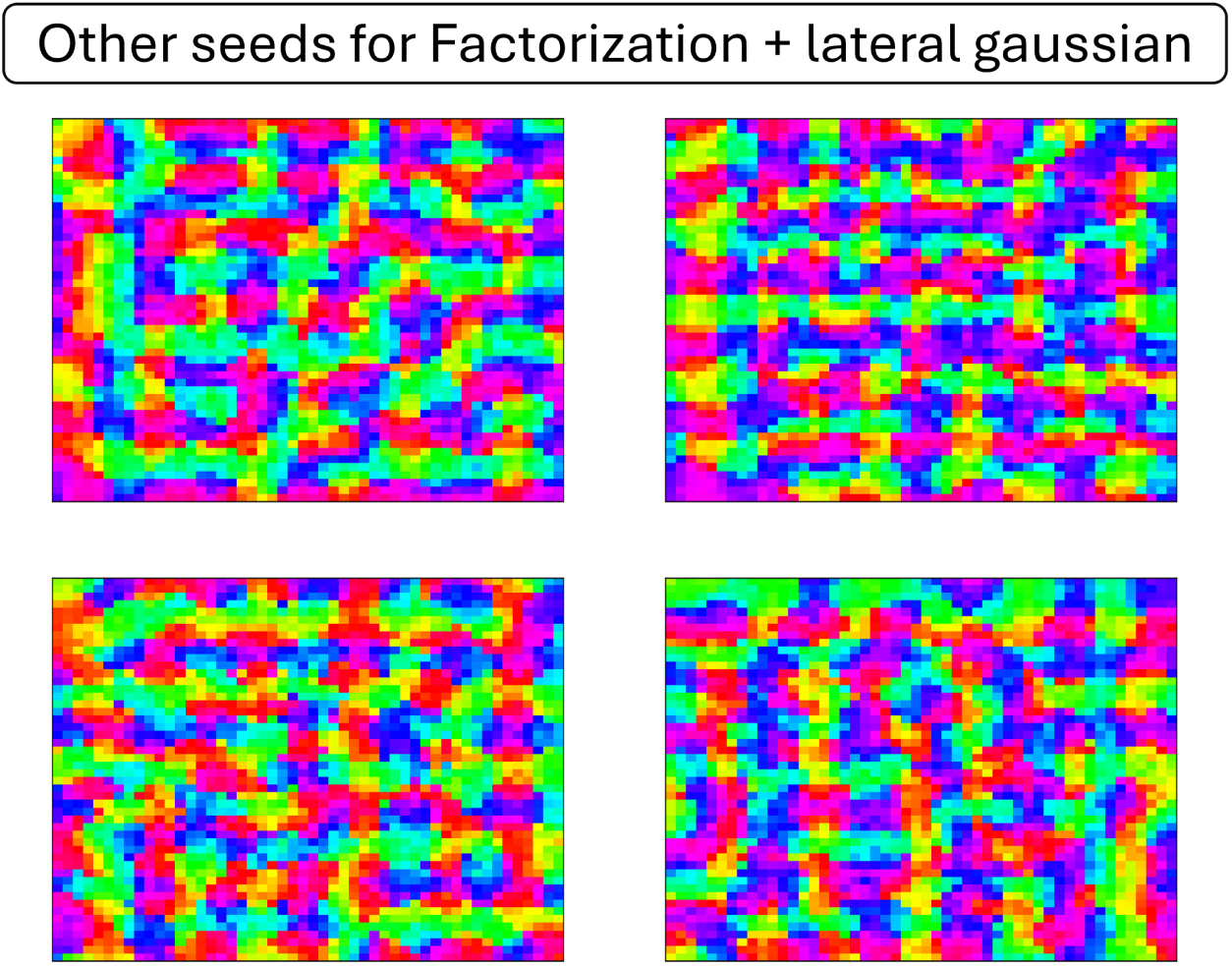
Other seeds for the factorization + Gaussian case.

**Fig. S8:**
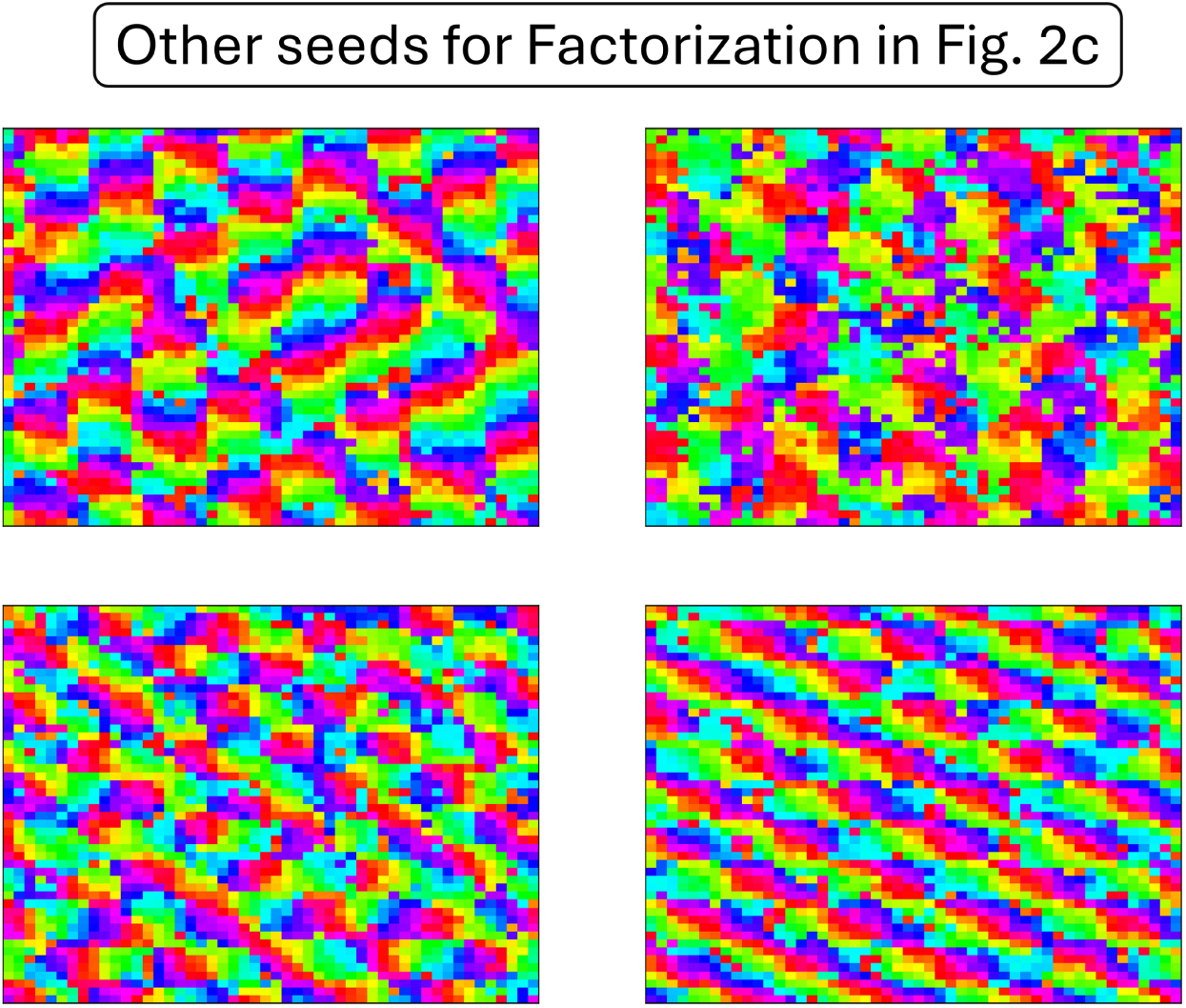
Other seeds for the Factorization case in. **Fig. 2c** **in the main text.**

**Fig. S9:**
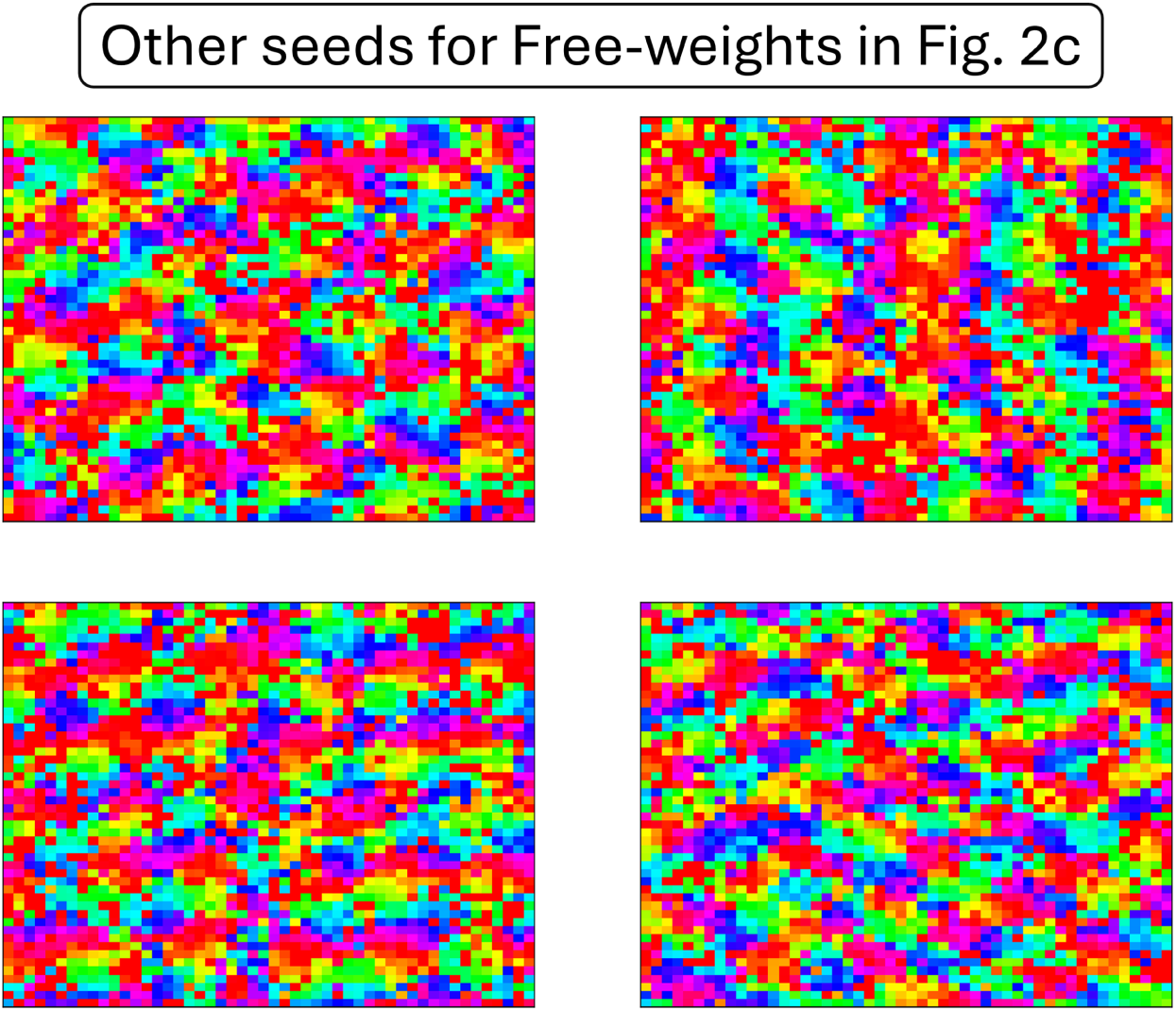
Other seeds for the Free-weights case in. **Fig. 2c** **in the main text.**

**Fig. S10:**
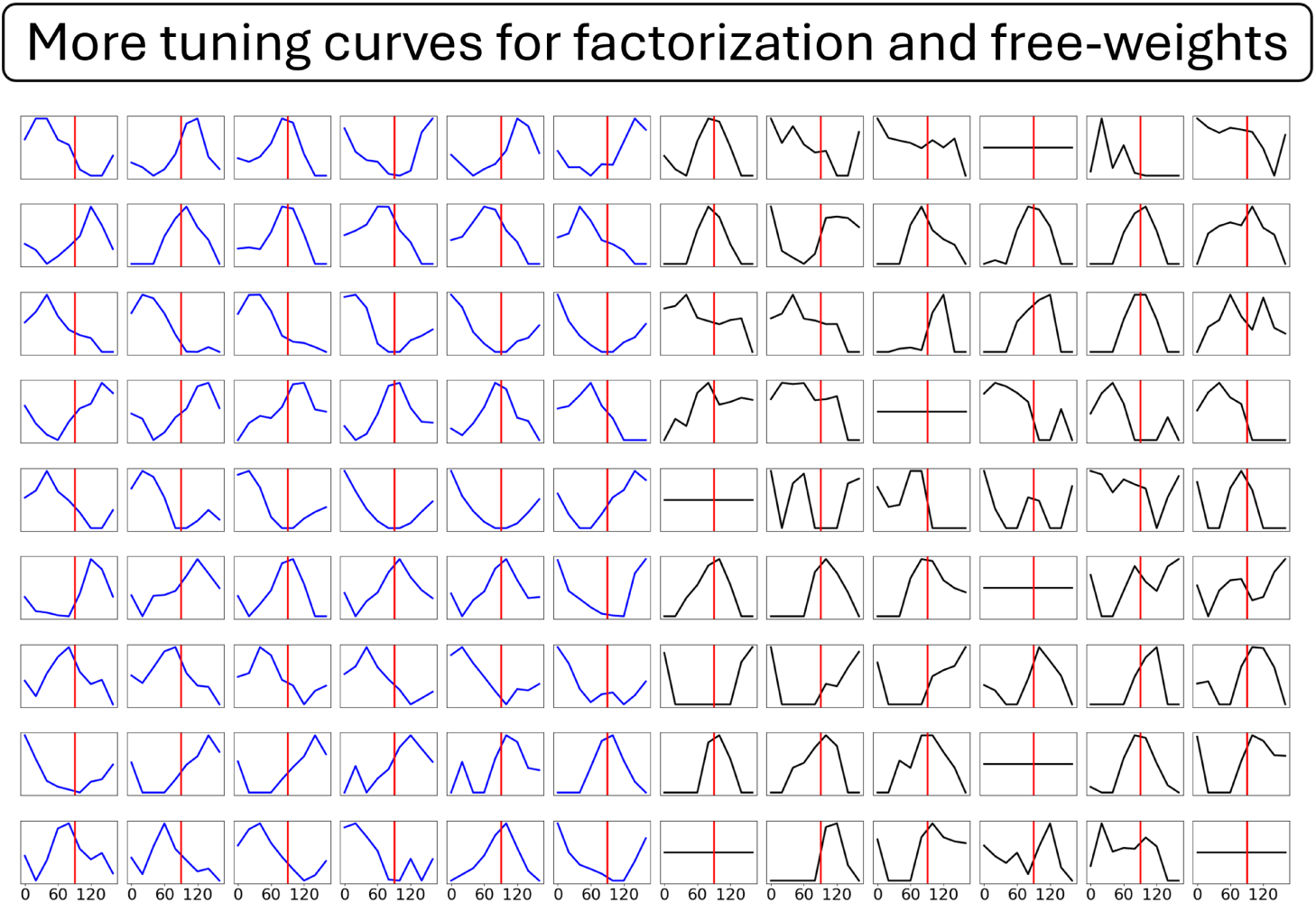
Other tuning curves for the neurons/maps shown in. **Fig. 2c** **of the main text. Blue is factorization, while black is free-weights.**

**Fig. S11:**
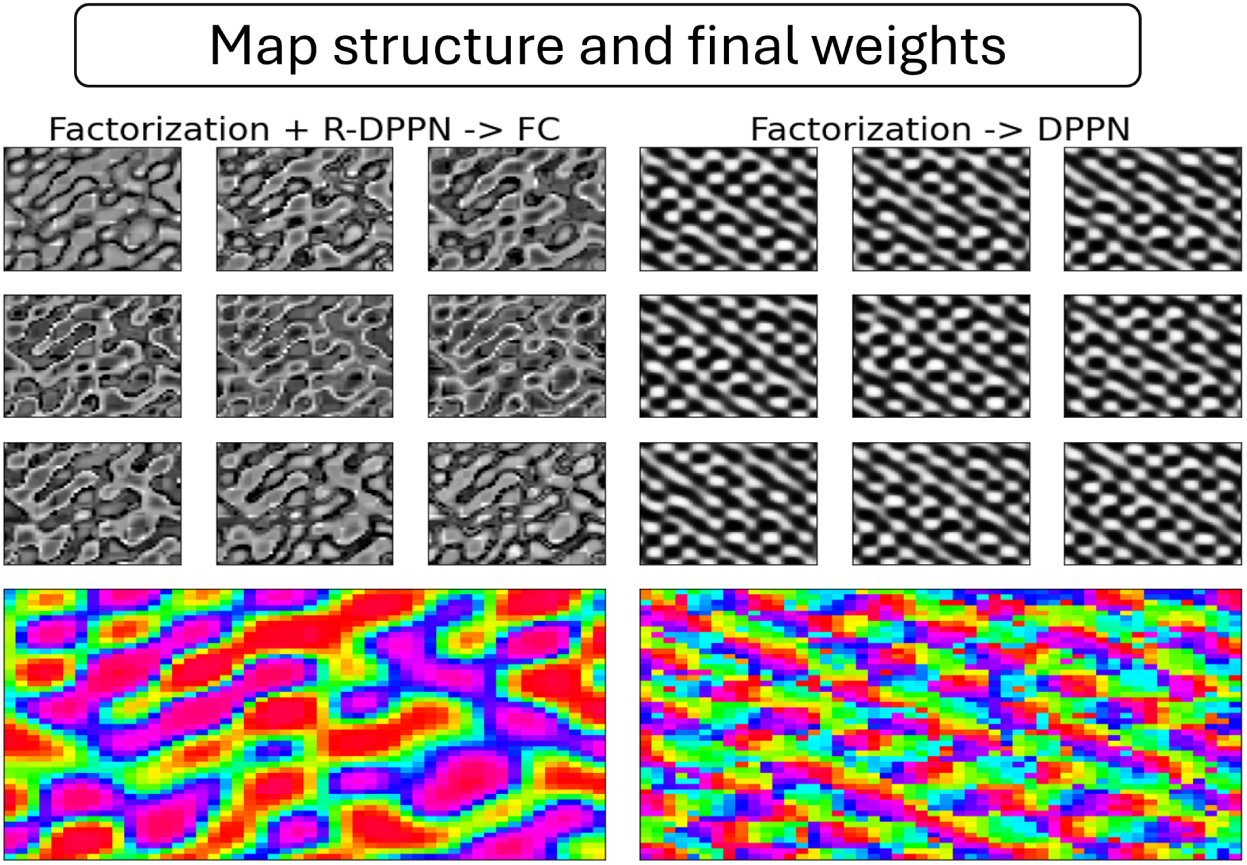
Correlation between the map structure and the final weights. This is done for two cases, where i) continuity is applied on the hidden layer by a R-DPPN (left image), or continuity is applied by the DPPN in the final layer (right images).

**Fig. S12:**
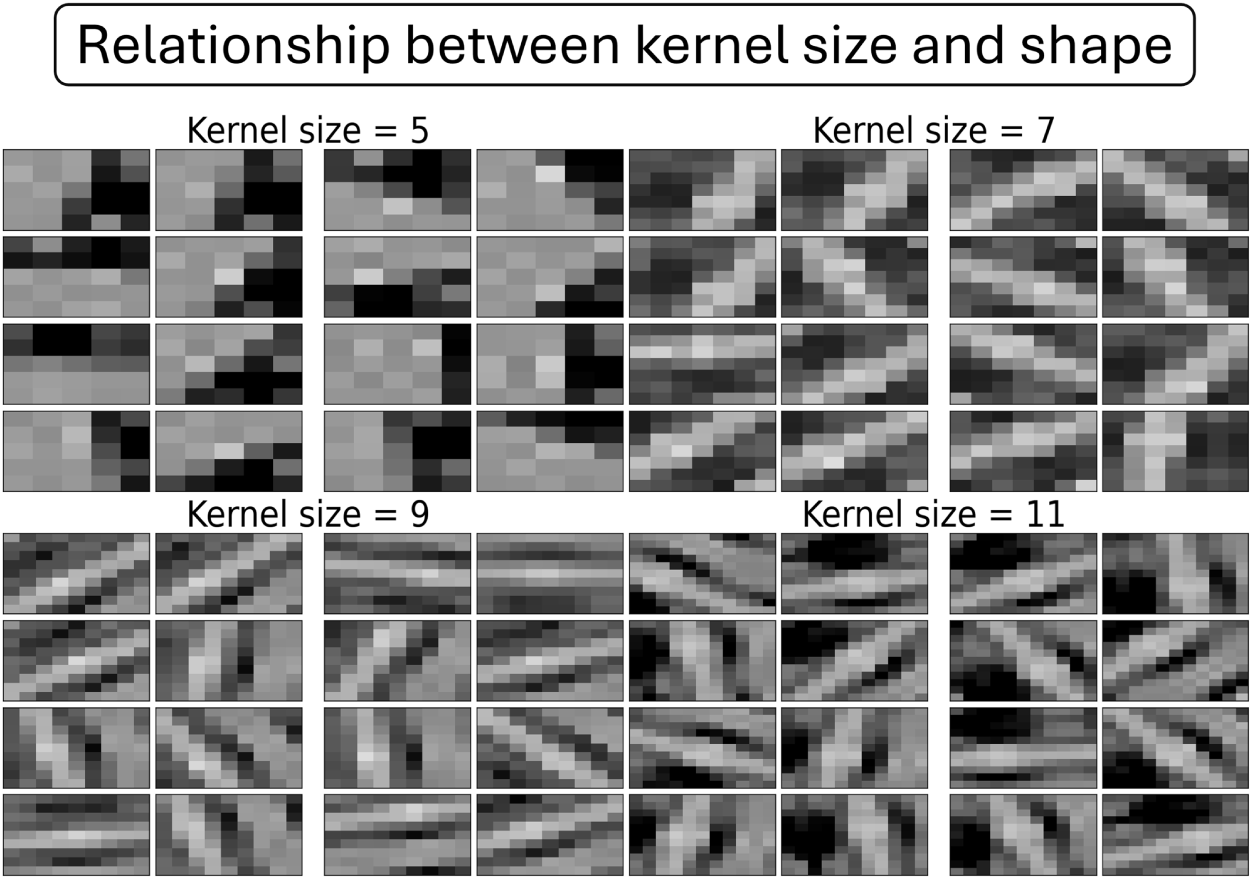
**Relationship between kernel size and kernel shape.**

**Fig. S13:**
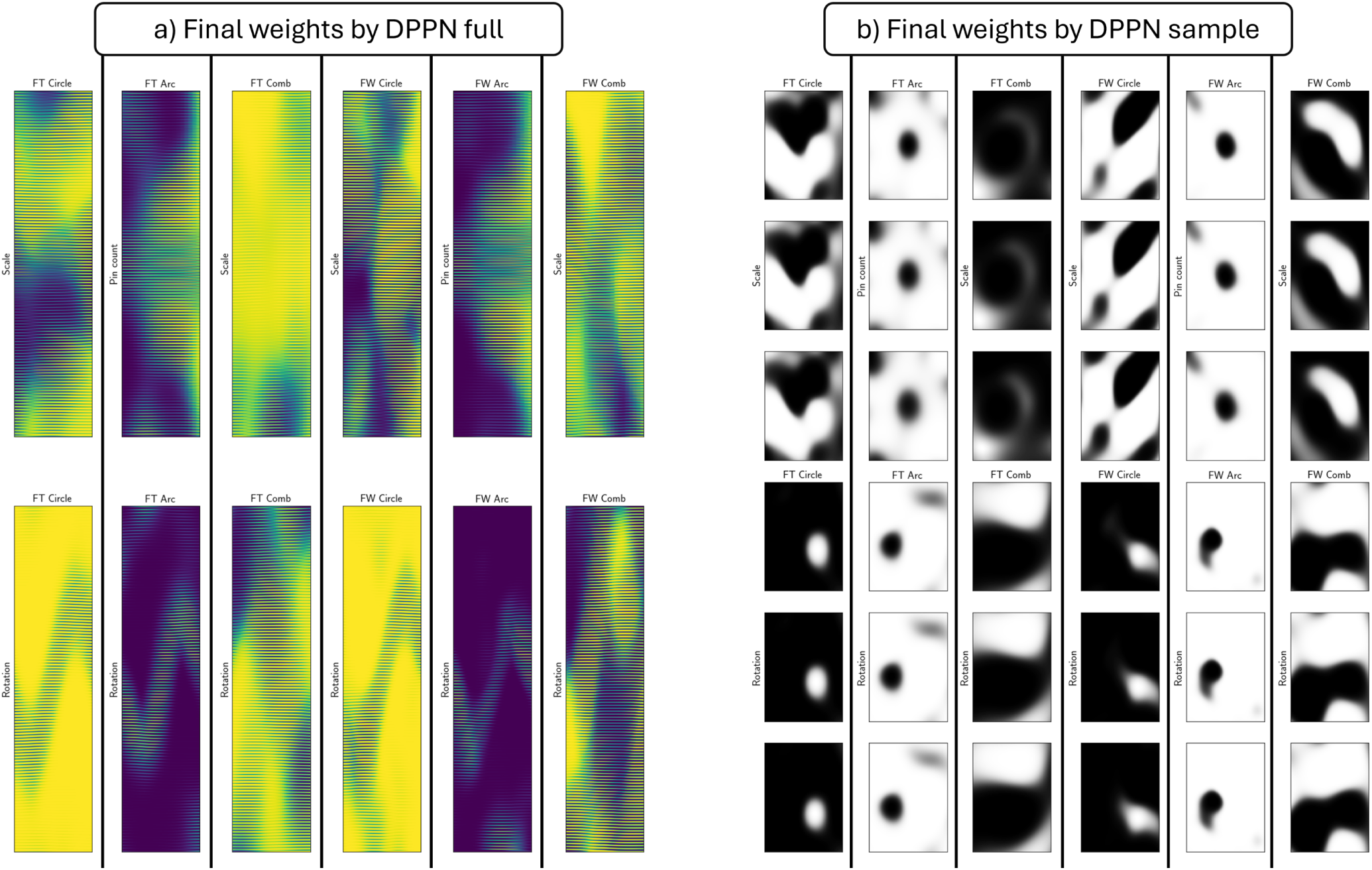
Phenotypic final weights as learned by the DPPN. a) full DPPN weights for each problem type Circle, Arc, Comb, and hidden layer architectures Factorization (FT) and free-weights (FW). b) the same configuration as a), but showing three sample weights for each problem and architecture types. The weights are chosen in succession (nearby in the layer).

**Fig. S14:**
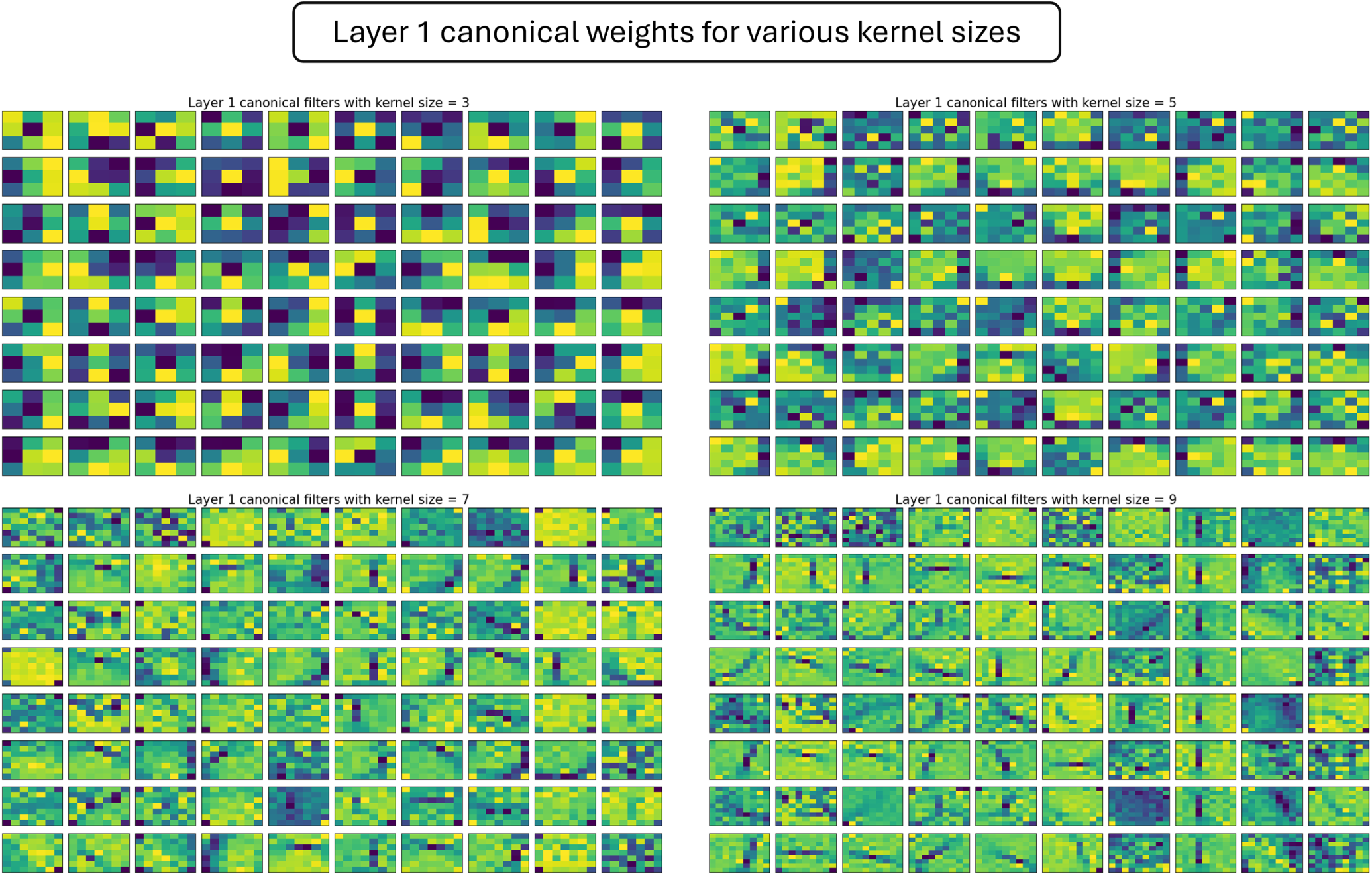
**Weights of efficient coding showing the canonical weights learned for the first layer with kernel sizes ∈ {3, 5, 7, 9}.**

**Table E1:**
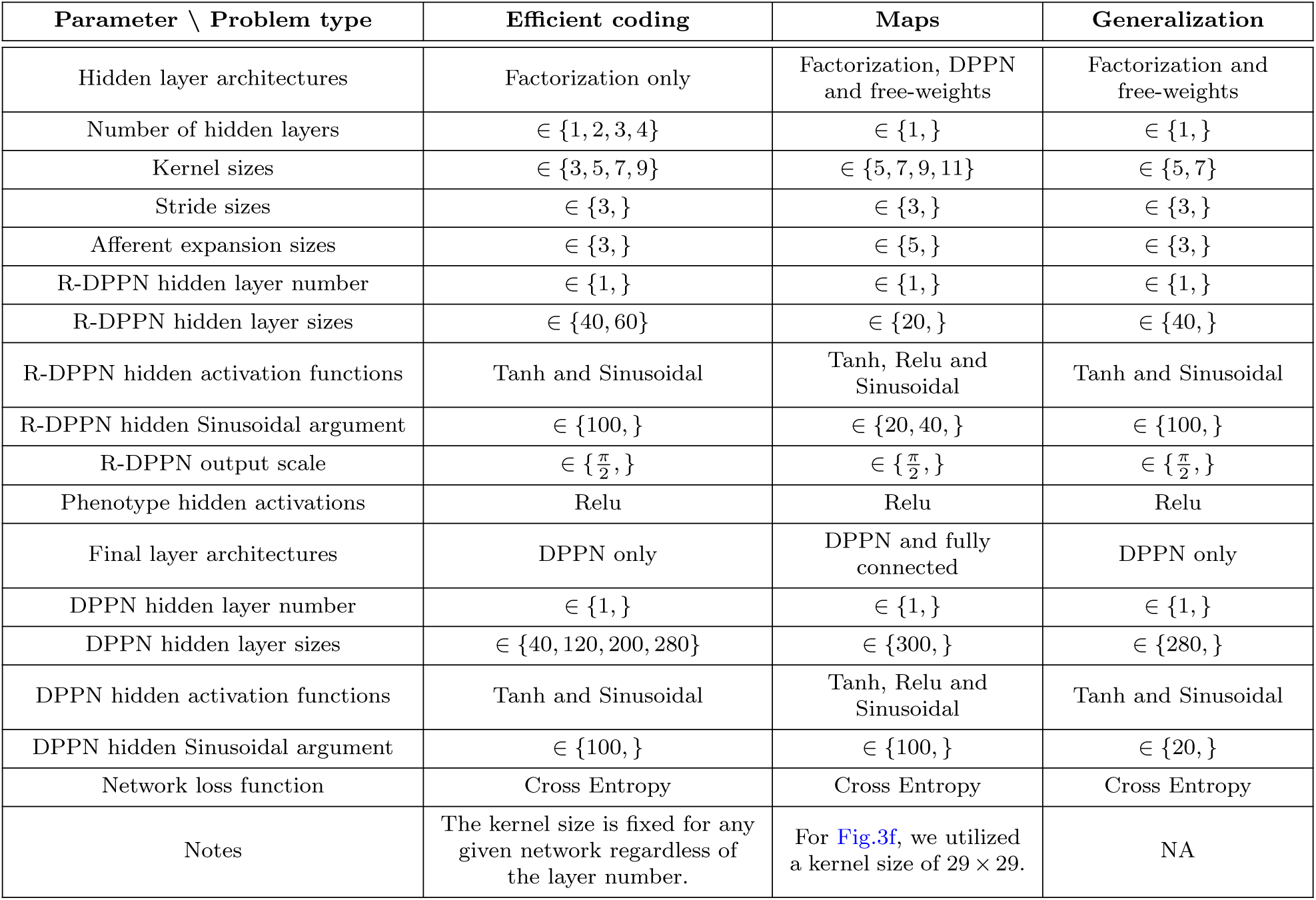
Network parameters.

**Table S1:** Summary of DPPN Vs. Factorization + R-DPPN for map generation.

|  | Architecture | Parameter count | Performance | Convergence to 0.3 loss | Directionality |
| --- | --- | --- | --- | --- | --- |
| Hidden size 1 | DPPN | 51 | $0.27 \pm 0.07$ | $2534 \pm 381$ | $0.49 \pm 0.13$ |
| | Factorization + R-DPPN | 52 | $0.11 \pm 0.12$ | $812 \pm 295$ | $0.80 \pm 0.11$ |
| Hidden size 2 | DPPN | 102 | $0.08 \pm 0.03$ | $985 \pm 116$ | $0.72 \pm 0.10$ |
| | Factorization + R-DPPN | 55 | $0.08 \pm 0.10$ | $626 \pm 42$ | $0.82 \pm 0.11$ |
| Hidden size 3 | DPPN | 153 | $0.05 \pm 0.02$ | $854 \pm 77$ | $0.62 \pm 0.16$ |
| | Factorization + R-DPPN | 58 | $0.06 \pm 0.07$ | $670 \pm 53$ | $0.84 \pm 0.08$ |
| Hidden size 4 | DPPN | 204 | $0.06 \pm 0.02$ | $858 \pm 56$ | $0.63 \pm 0.15$ |
| | Factorization + R-DPPN | 61 | $0.04 \pm 0.01$ | $687 \pm 52$ | $0.83 \pm 0.09$ |
| Hidden size 5 | DPPN | 255 | $0.06 \pm 0.02$ | $719 \pm 23$ | $0.61 \pm 0.16$ |
| | Factorization + R-DPPN | 64 | $0.05 \pm 0.02$ | $695 \pm 33$ | $0.84 \pm 0.09$ |
| Hidden size 6 | DPPN | 210 | $0.07 \pm 0.02$ | $819 \pm 14$ | $0.62 \pm 0.15$ |
| | Factorization + R-DPPN | 79 | $0.05 \pm 0.01$ | $725 \pm 12$ | $0.88 \pm 0.09$ |
| Hidden size 7 | DPPN | 1020 | $0.09 \pm 0.03$ | $788 \pm 17$ | $0.63 \pm 0.15$ |
| | Factorization + R-DPPN | 104 | $0.06 \pm 0.02$ | $780 \pm 48$ | $0.85 \pm 0.09$ |

## References

1. Arnqvist, G., Sayadi, A., Immonen, E., Hotzy, C., Rankin, D., Tuda, M., . . . Johnston, J. S. (2015). Genome size correlates with reproductive fitness in seed beetles. *Proc. Biol. Sci*., *282*, 20151421. Retrieved from 10.1098/rspb.2015.1421

2. Bednar, J. A. (2012). Building a mechanistic model of the development and function of the primary visual cortex. J. Physiol. Paris., 106, 194–211. Retrieved from 10.1016/ j.jphysparis.2011.12.001

3. Bernardi, S., Benna, M. K., Rigotti, M., Munuera, J., Fusi, S., & Salzman, C. D. (2020). The geometry of abstraction in the hippocampus and prefrontal cortex. Cell, 183, 954-67.e21. Retrieved from 10.1016/j.cell.2020.09.031

4. Blasdel, G. G., & Salama, G. (1986). Voltage-sensitive dyes reveal a modular organization in monkey striate cortex. Nature, 321, 579–85. Retrieved from 10.1038/321579a0

5. Cloherty, S. L., Hughes, N. J., Hietanen, M. A., Bhagavatula, P. S., Goodhill, G. J., & Ibbotson, M. R. (2016). Sensory experience modifies feature map relationships in visual cortex. Elife, 5, e13911. Retrieved from 10.7554/eLife.13911

6. Clune, J., Stanley, K. O., Pennock, R. T., & Ofria, C. (2011). On the performance of indirect encoding across the continuum of regularity. IEEE Trans. Evol. Comput., 15, 346-67. Retrieved from 10.1109/TEVC.2010.2104157

7. Crair, M. C., Gillespie, D. C., & Stryker, M. P. (1998). The role of visual experience in the development of columns in cat visual cortex. Science, 279, 566-70. Retrieved from https:// doi.org/10.1126/science.279.5350.566

8. Crowley, J., & Katz, J. (1999). Development of ocular dominance columns in the absence of retinal input. Nat. Neurosci., 2, 1125–30. Retrieved from 10.1038/16051

9. Durbin, R., & Mitchison, G. (1990). A dimension reduction framework for understanding cortical maps. Nature, 343, 644–7. Retrieved from 10.1038/343644a0

10. Fernando, C., Banarse, D., Reynolds, M., Besse, F., Pfau, D., Jaderberg, M., . . . Wierstra, D. (2016). Convolution by evolution: Differentiable pattern producing networks. GECCO ’16: Proceedings of the Genetic and Evolutionary Computation Conference 2016, 109–16. Retrieved from 10.1145/2908812.2908890

11. Ghirlanda, S., & Enquist, M. (2003). A century of generalization. Animal Behaviour, 66, 15-36. Retrieved from 10.1006/anbe.2003.2174

12. Gunasekar, S., Woodworth, B., Bhojanapalli, S., Neyshabur, B., & Srebro, N. (2017). Implicit regularization in matrix factorization. arXiv . Retrieved from 10.48550/ arXiv.1705.09280

13. Ha, D., Dai, A., & Le, Q. V. (2016). Hypernetworks. arXiv . Retrieved from 10.48550/arXiv.1609.09106

14. Han, Y., Roig, G., Geiger, G., & Poggio, T. (2020). Scale and translation-invariance for novel objects in human vision. Sci. Rep., 10, 1411. Retrieved from 10.1038/s41598-019-57261-6

15. Hu, E. J., Shen, Y., Wallis, P., Allen-Zhu, Z., Li, Y., Wang, S., . . . Chen, W. (2021). Lora: Low-rank adaptation of large language models. arXiv . Retrieved from 10.48550/ arXiv.2106.09685

16. Hulse-Kemp, A. M., Maheshwari, S., Stoffel, K., Hill, T. A., Jaffe, D., Williams, S. R., . . . Deynze, A. V. (2018). Reference quality assembly of the 3.5-gb genome of capsicum annuum from a single linked-read library. Hortic. Res., 5. Retrieved from 10.1038/s41438-017-0011-0

17. Jaderberg, M., Simonyan, K., Zisserman, A., & Kavukcuoglu, K. (2015). Spatial transformer networks. Advances in neural information processing systems, 28. Retrieved from 10.48550/arXiv.1506.02025

18. Kaschube, M., Schnabel, M., Lowel, S., Coppola, D. M., White, L. E., & Wolf, F. (2010). Universality in the evolution of orientation columns in the visual cortex. Science, 330, 1113–16. Retrieved from 10.1126/science.1194869

19. Kavukcuoglu, K., Sermanet, P., Boureau, Y., Gregor, K., Mathieu, M., & Cun, Y. (2010). Learning convolutional feature hierarchies for visual recognition. NeurIPS, 23.

20. Kobylkov, D., Rosa-Salva, O., Zanon, M., & Vallortigara, G. (2024). Innate face-selectivity in the brain of young domestic chicks. PNAS, 121, e2410404121. Retrieved from 10.1073/pnas.2410404121

21. LeCun, Y., Bottou, L., Bengio, Y., & Haffner, P. (2002). Gradient-based learning applied to document recognition. Proceedings of the IEEE, 86, 2278–324. Retrieved from 10.1109/5.726791

22. Malerba, M. E., Ghedini, G., & Marshall, D. J. (2020). Genome size affects fitness in the eukaryotic alga dunaliella tertiolecta. Curr. Biol., 30, 3450–6.e3. Retrieved from 10.1016/j.cub.2020.06.033

23. Novikov, A., Podoprikhin, D., Osokin, A., & Vetrov, D. (2015). Tensorizing neural networks. arXiv. Retrieved from 10.48550/arXiv.1509.06569

24. O’Keefe, J., & Dostrovsky, J. (1971). The hippocampus as a spatial map. preliminary evidence from unit activity in the freely-moving rat. Brain Res., 34, 171–5. Retrieved from 10.1016/0006-8993(71)90358-1

25. Olshausen, B. A., & Field, D. J. (1997). Sparse coding with an overcomplete basis set: A strategy employed by v1? Vision Research, 37, 3311–25. Retrieved from 10.1016/S0042-6989(97)00169-7

26. Philips, R. T., & Chakravarthy, V. S. (2017). A global orientation map in the primary visual cortex (v1): Could a self organizing model reveal its hidden bias? Front. Neural. Circuits, 10, 109. Retrieved from 10.3389/fncir.2016.00109

27. Qian, X., Dehghani, A. O., Farahani, A. B., & Bashivan, P. (2026). Local lateral connectivity is sufficient for replicating cortex-like topographical organization in deep neural networks. Nat. Commun., 17, 4042. Retrieved from 10.1038/s41467-026-70065-3

28. Salva, O. R., Farroni, T., Regolin, L., Vallortigara, G., & Johnson, M. H. (2011). The evolution of social orienting: Evidence from chicks (gallus gallus) and human newborns. PloS one, 6, e18802. Retrieved from 10.1371/journal.pone.0018802

29. Shuvaeva, S., Lachia, D., Koulakova, A., & Zador, A. (2024). Encoding innate ability through a genomic bottleneck. PNAS, 121, e2409160121. Retrieved from 10.1073/pnas.2409160121

30. Sosa, F. A., & Stanley, K. O. (2018). Deep hyperneat : Evolving the size and depth of the substrate evolutionary.. Retrieved from https://api.semanticscholar.org/CorpusID:54025867

31. Stanley, K. O. (2007). Compositional pattern producing networks: A novel abstraction of development. Genet. Program. Evolvable Mach., 8, 131–62. Retrieved from 10.1007/s10710-007-9028-8

32. Stanley, K. O., D’Ambrosio, D. B., & Gauci, J. (2009). A hypercube-based encoding for evolving large-scale neural networks. Artificial life, 15, 185–212. Retrieved from 10.1162/artl.2009.15.2.15202

33. Stevens, J.-L. R., Law, J. S., Antolık, J., & Bednar, J. A. (2013). Mechanisms for stable, robust, and adaptive development of orientation maps in the primary visual cortex. J. Neurosci., 33, 15747–66. Retrieved from 10.1523/JNEUROSCI.1037-13.2013

34. Su, J., Vargas, D. V., & Sakurai, K. (2019). One pixel attack for fooling deep neural networks. IEEE Trans. Evol. Comput., 23, 828–41. Retrieved from 10.1109/TEVC.2019.2890858

35. Swaminathan, S., Garg, D., Kannan, R., & Andres, F. (2020). Sparse low-rank factorization for deep neural network compression. Neurocomputing, 398, 185–96. Retrieved from 10.1016/j.neucom.2020.02.035

36. Teichert, T., Wachtler, T., Michler, F., Gail, A., & Eckhorn, R. (2007). Scale-invariance of receptive field properties in primary visual cortex. BMC Neurosci., 8, 38. Retrieved from 10.1186/1471-2202-8-38

37. Villalba, L. S., Calangiu, I., Boehringer, R., Mante, V., & Grewe, B. F. (2025). Category learning disentangles representation of trial events in hippocampus ca1. bioRxiv. Retrieved from 10.64898/2025.12.01.690977

38. Wang, S., Vasas, V., Freeland, L., Osorio, D., & Versace, E. (2024). Spontaneous biases enhance generalization in the neonate brain. Iscience, 27, 110195. Retrieved from 10.1016/j.isci.2024.110195

39. White, L., D., D. C., & Fitzpatrick, D. (2001). The contribution of sensory experience to the maturation of orientation selectivity in ferret visual cortex. Nature, 411, 1049–52. Retrieved from 10.1038/35082568

40. Wiecek, E., Ramirez, L. D., Klimova, M., & Ling, S. (2026). Spatial frequency tuning follows scale invariance in the human visual cortex. J. Neurosci., 46, e1490252025. Retrieved from 10.1523/JNEUROSCI.1490-25.2025

